# Beyond Pollination: Computer Vision Reveals how Flower Visitors, Climate and Agroforestry Management Drive Cocoa Yields in China and Brazil

**DOI:** 10.1101/2025.10.07.680934

**Authors:** Manuel Toledo-Hernández, Marcel Balle, Julian Barillaro, Kevin F. A. Darras, Valentina Fortunato, Fupeng Li, Mirco Solé, Xiaoye Tong, Wenxiu Xu, Thomas C. Wanger

## Abstract

Cocoa (*Theobroma cacao* L.) is a multi-billion-dollar crop that is strongly affected by climate change. As a pollination-limited crop, improving little-understood pollination services in agroforestry systems may offer a scalable solution to increase yield sustainably in a changing climate. Here, we use embedded computer vision devices and structural equation models to quantify the interactions of flower visitors and temperature across shade tree-diversity and canopy cover gradients in cocoa systems on fruit set, as a yield precursor. In China and Brazil we show that flower visits are done by nectar and pollen foragers (23.1%), herbivores (6.1%), predators (2.3%), and visitors combining the three functions (63%). Forager visits were driven by increased shade-tree diversity and canopy cover management, with stronger effects in China than in Brazil. Foraging midges in China and multifunctional ants in Brazil enhanced fruit set, showing that diverse pollinators across continents affect cocoa yields. In China, higher canopy cover reduced aphids foraging and feeding on flower tissue, while in Brazil temperature increase reduced flowering. Overall, new technologies can guide implementation of agroforestry management strategies to enhance pollination while reducing pest pressures to ensure sustainable cocoa production under climate change.

## Introduction

Chocolate, a globally cherished product derived from the beans of the cocoa tree (*Theobroma cacao* L.), is a USD 140 billion industry facing significant threats from climate change (Kongor et al., 2024). In 2024, abnormal temperature and rainfall patterns in Ghana and Ivory Coast, the world’s leading cocoa producers, caused a 24% bean production decline compared to the previous year (ICCO, 2024a,b). This supply deficit tripled cocoa bean prices within the same year (Trading Economics, 2025) and rose concerns for massive cocoa production expansion into primary forest areas (GFW, 2025) as well as supply chain issues for the livelihoods of 5 million smallholder farmers (Beg et a., 2017). Rising cocoa prices also made cocoa production in other areas such as China and Brazil more attractive. In China, the expansion of newly introduced cocoa production is planned (Li et al. 2022) and in Brazil, cocoa production is encouraged after the sector is recovering from decades of pest and disease outbreaks (Marques et al., 2021). Tackling these climate-driven socio-environmental challenges will require a fundamental shift towards sustainable cocoa production that secures cocoa supplies while protecting people’s livelihoods and nature.

Impact investments into agroforestry approaches have been promoted to maintain ecosystem services such as soil fertility, carbon sequestration potential and pollination (Tscharntke et al., 2011; ICI, 2025; WCF, 2025). By sustaining these ecosystem services, agroforestry can not only enhance agricultural production, but it can also prevent forest conversion, thereby supporting compliance with international zero-deforestation regulations (Gilbert, 2024). While a critical argument against cocoa agroforest are lower yields compared to monocultures (Niether et al. 2020), hand pollination can enhance yields in Indonesia and Brazil by up to 700% in this pollination limited crop (Toledo-Hernández et al., 2020; 2023). Agroforestry management can buffer yield losses that results from pollination deficits and climate change through an appropriate selection of shade tree species and canopy cover adjustments that both buffer temperature increases and enhance pollinator habitats (Lander et al., 2025; Toledo-Hernández et al., 2017).

Currently, however, most work on cocoa pollination relies on hand pollination as a surrogate for the maximum yield potential of a fully pollinated tree. The identity of pollinators, both locally and globally, remains unknown, as does their response to plantation management practices, including pesticide use. A major research challenge remains to identify effective pollinators among the large diversity of small flower visitors and a cocoa flower opening time of only 48-hours to observe pollination (Toledo-Hernández et al., 2017). New biodiversity monitoring technologies such as computer vision (Høye et al., 2021; Bjerge et a., 2022; Darras et al., 2024) offer promising, yet untapped opportunities in cocoa production for real-time and accurate observations of small flower visitors, their ecological functions and the effects on cocoa yields.

Here, we aim to quantify the effects of cocoa flower visitors and climate change on yields with embedded computer vision (ECV) cameras at 21 research plots across a shade tree diversity and canopy cover gradient in cocoa systems of China and Brazil. We (i) identify cocoa flower visitors and their ecological functions, to (ii) evaluate the role of shade tree species richness and canopy cover in supporting these functions. We then combine our data with historical spatiotemporal temperature datasets (Muñoz Sabater, 2019) to (iii) disentangle the complex interactions among climate (temperature), agroforestry management (shade tree-diversity and canopy cover), and main flower visitors driving fruit set as a yield precursor.

## Results & Discussion

### Cocoa flower visitors and their ecological functions

With our ECV cameras (Darras et al., 2024), we monitored 1,071 individual cocoa flowers in 128 cocoa trees, distributed in 21 research plots from March to July 2023 in China and from May to August 2023 in Brazil (Supplementary Table S1, S2). Across 23,414 hours of continuous observations, the cameras detected visitors in 31.2% of all flowers in China and 80.1% of all flowers in Brazil. The ECV cameras showed 1.4 visitors/flower in China and 11.5 visitors/flower in Brazil, which exceeds documented visitation rates in Indonesia (0.1 visitors/flower; Toledo-Hernández et al., 2021), Peru (0.2 visitors/flower; Vansynghel et al., 2022), and Bolivia (0.3 visitors/flower; Chumacero de Schawe et al., 2018) monitored with adhesive traps. This suggests that our ECV camera outperforms traditional methods in detecting tiny arthropods and their real number of visits in cocoa flowers. A limitation of adhesive traps may be that the glue blocks aerial chemical attractants to flower visitors, thereby reducing flower visits. We registered 4,766 single visits belonging to 12 taxonomic groups and 30 morpho-species in China, and 15 taxonomic groups and 42 morpho-species in Brazil (Fig. 1a,b; Table 1, Supplementary Figure S1, S2, Table S3, S4). Overall, this confirms previous monitoring evidence with yellow pan traps, adhesive traps on flowers, and photos and video recordings that cocoa flowers are visited by a diverse range of arthropods (Toledo-Hernández et al., 2017; Claus et al., 2018; Vansynghel et al., 2022; Jordao et al., 2024) but also suggests that visitation rates are substantially underestimated.

**Figure 1.**
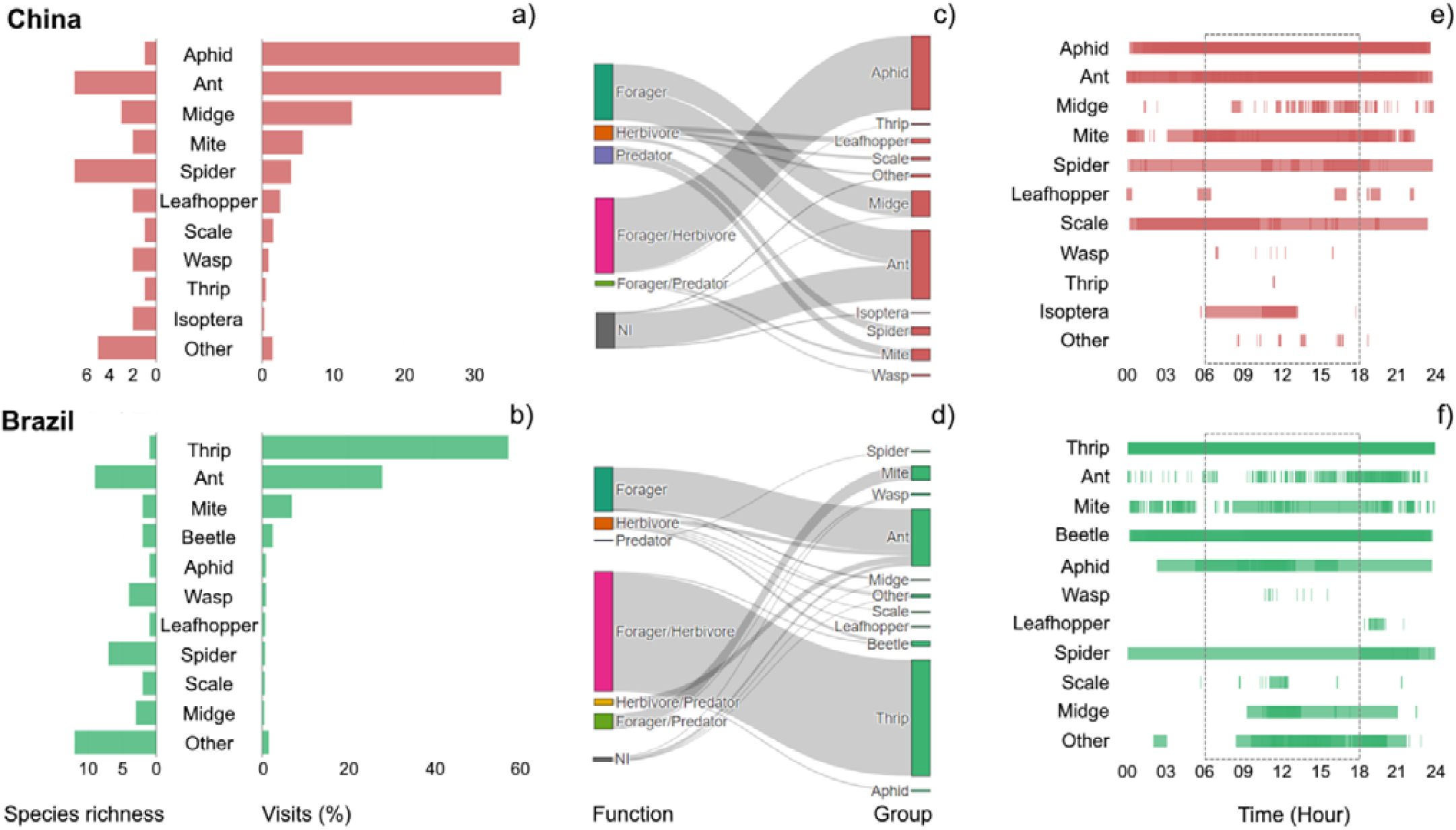
Cocoa flower visitor communities in China and Brazil. In Hainan, China, we recorded 12 taxonomic groups. Aphids, ants, and midges were the most frequent visitors, and ants, spiders and midges were the most diversified (a). In Bahia, Brazil, we identified 15 taxonomic groups, with thrip, ant, and mite as the most frequent visitors, and ants, spiders, and wasps the most diversified (b). We classify visitors based on the presence and ecological functions based on their behavior in flowers (c, d). Visitor functions are forager (i.e. nectar and/or pollen feeder), herbivore (i.e. flower sap and/or tissue feeder), predator (i.e. predating and/or scouting prey), and the combinations of these three functions. We recorded the cumulative duration of visits per flower visitor group over a 24-hour period (e, f). Darker colors indicate overlapping visits by multiple visitors, and daytime visitation (06:00–18:00) is highlighted with a grey dashed box. The group “others” represented beetles, and stinkbugs in China; and flies, mealybugs, bees, grasshoppers and isopterans in Brazil. NI in panel “c” and “d” stands for “not identified” function.

**Table 1.**
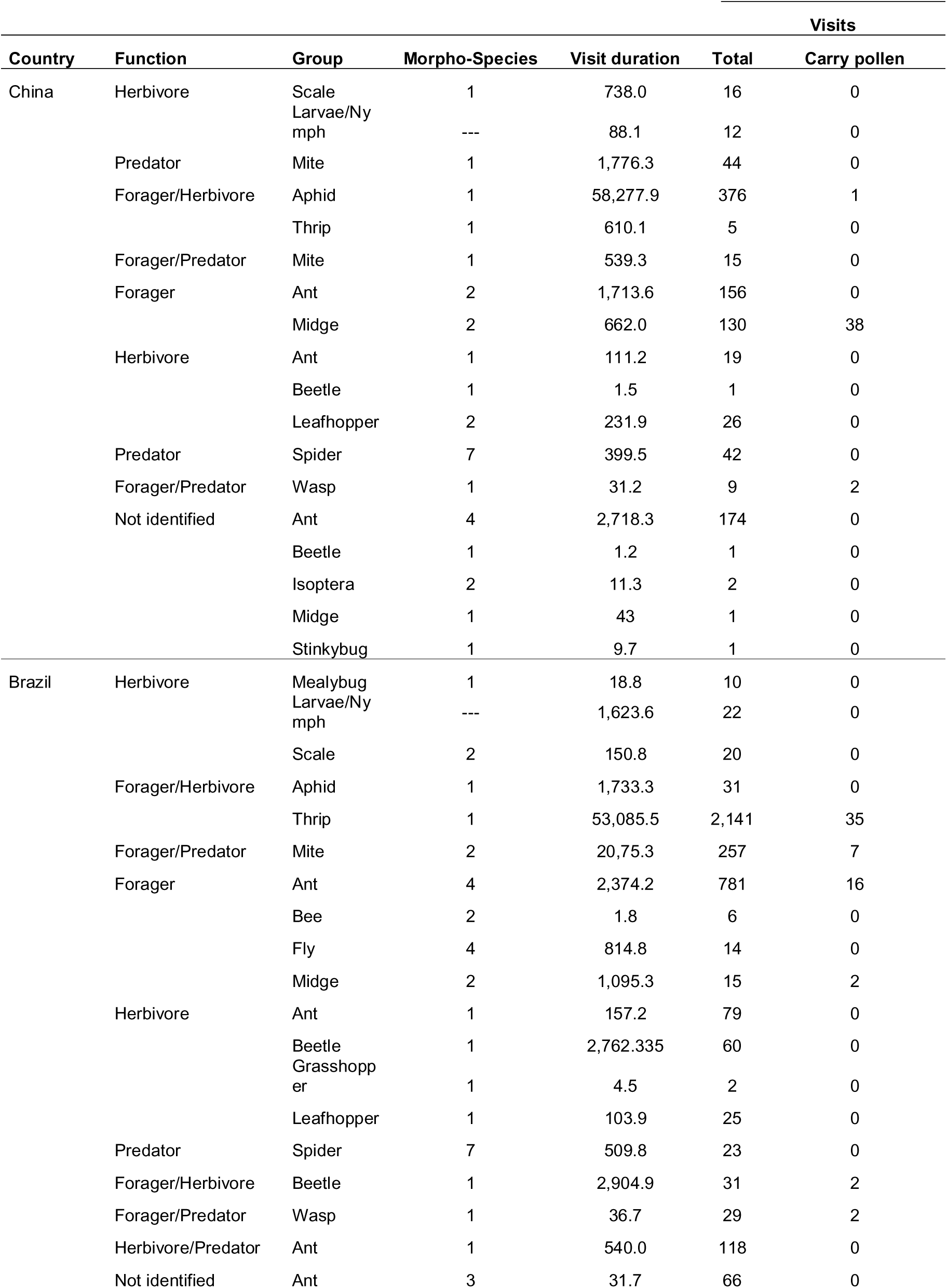
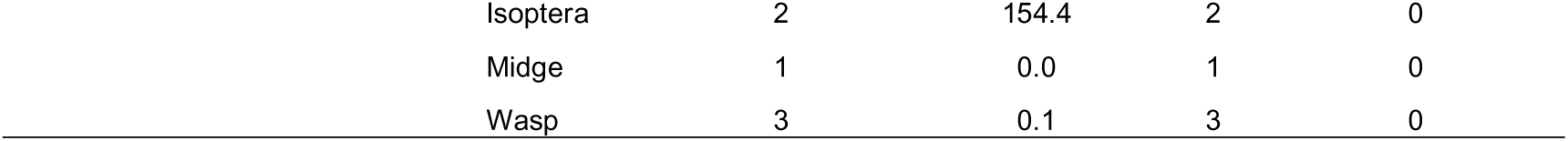
Summary of cocoa flower visitors in China and Brazil. Visitors are categorized based on their three basic ecological functions (i.e. foragers or feeding on pollen and/or nectar, herbivores or feeding on flower sap or tissue, predators or feeding on or scouting prey) and the combination of them (i.e. forager/herbivore, forager/predator, herbivore/predator). Visit duration is the cumulative minutes recorded in flowers. Visits refer to the number of visits recorded. Carry pollen stands for visits with visible pollen in visitor body.

The macro-photography and 24h (day and night) continuous observations capabilities of our ECV cameras allowed us to record flower visitor behavior in different flower structures (i.e. petals, anthers, style, and staminodes), and to determine their ecological functions based on their observed behavior and time spent in flowers (Fig. 1c-f, Supplementary Figure S3, Table S5). We recorded 23.1% of total visitors feeding on pollen and nectar (i.e. foragers), 6.1% were flower-herbivores piercing and cutting floral tissue, 2.3% were predators actively scouting for prey; and 63% of visitors had a combination of the three ecological functions (Fig. 1c,d). Seven taxonomic groups were observed in the images transporting pollen on their bodies, primarily midges (China = 29.2% of visits; Brazil = 13.3% of visits) and parasitoid wasps (China = 22.2% of visits; Brazil = 6.9% of visits; all pollen transporting visitors are in Table. 1). When considering the cumulative time spent in flowers, aphids (85.7 %), ants (6.7 %) and mites (2.6 %) were the most important groups in China, and thrips (75.6 %), beetles (8.1 %) and ants (4.4 %) in Brazil (Fig. 1e,f, Table 1). These findings suggest that contrary to the widely accepted single-pollinator hypothesis (Toledo-Hernández et al., 2017; Claus et al., 2018), pollination success is likely dependent on multiple plant-arthropod interactions.

### Agroforests mediate flower foragers and herbivores

In Hainan, China, most cocoa areas are cultivated in unshaded monocultures and to a lesser extent in agroforests with single palm tree species such as areca and coconut (Zhang et al., 2021; Supplementary Figure S4a). In Bahia, Brazil, on the contrary, cocoa is integrated into the Atlantic Forest as diverse agroforestry systems known as cabrucas (Cassano et al., 2012; Supplementary Figure S4b). To quantify the effect of shade-tree species diversity and canopy cover on flower visitors in these cocoa systems, we split our 21 research plots into six distinctly managed agroforests following a gradient of increasing shade tree species richness and canopy cover; namely in China (i) monocultures (no shade trees), (ii) low (one shade palm species and <30% canopy cover) and (iii) intermediate (one shade palm species, and 40-50% canopy cover) shaded systems, and in Brazil (iv) low (1-3 tree species, and 30-40% canopy cover), (v) intermediate (4-5 tree species, and 40-50% canopy cover), and high (7-9 tree species, and >50% canopy cover) shaded systems; with three and four plot replicates per management category in China and Brazil, respectively (Supplementary Figure S4).

In China, our multiple comparison analysis showed significantly higher proportion of visits of nectar and pollen foragers in low (p=0.040), and intermediate (p=0.010) shaded systems compared to monocultures (Fig. 2a; Supplementary Table S6; S7). Combined forager/herbivore visitors were significantly higher in monocultures in contrast to systems with low (p=0.018) and intermediate (p=0.002) shade-tree management. In Brazil, foragers had a significantly higher proportion of visits in intermediate compared to low (p=0.035) and high (p=0.024) shaded systems. (Fig. 2b; Supplementary Table S6; S8). These results highlight the key role of intermediate shaded systems (i.e., shade cover between 40-50%) in conserving potential pollinators in both countries, and in the case of China, suppressing potential cocoa pests, like thrips and aphids, through shade tree species and canopy cover management (Vaast et al., 2014; Ambele et al., 2023). Moreover, in China, even a single shade tree species with canopy cover above 30% may support potential pollinators and help control flower herbivores. In contrast, enhancing potential pollinators in Brazil may require greater shade tree diversity and canopy cover exceeding 40%, highlighting the importance of understanding global variability in pollinator diversity and management.

**Figure 2.**
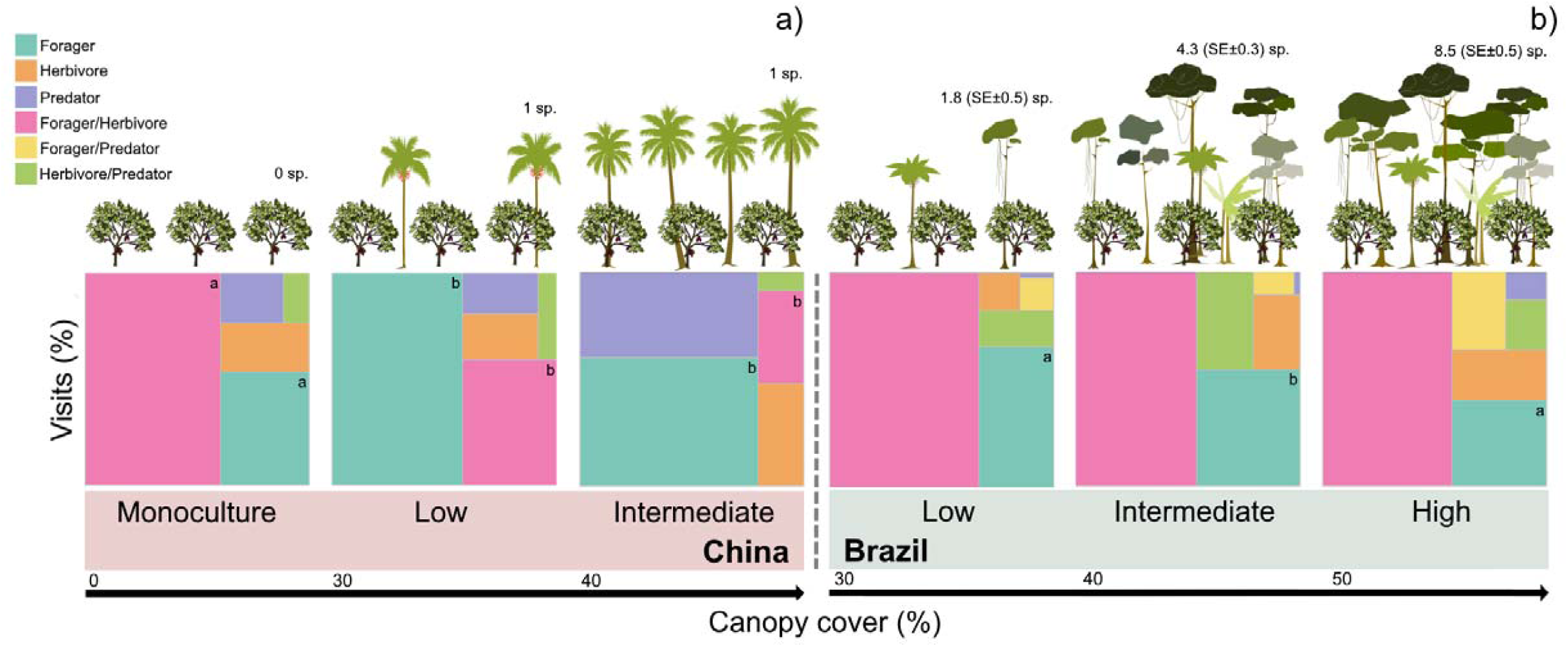
Ecological functions of cocoa flower visitors across agroforestry systems in China (a) and Brazil (b). Cocoa agroforestry systems were categorized based on shade-tree diversity and canopy cover. In China, categories were monoculture (no trees), low (one areca palm species, 30-40% canopy cover), and intermediate (one coconut palm species, 40-50% canopy cover). In Brazil, categories were low (1-3 tree species, 30-40% canopy cover), intermediate (4-5 tree species, 40-50% canopy cover), and high (7-9 tree species, >50% canopy cover). We assessed the proportion of visits of flower visitors performing different ecological functions: foragers (nectar/pollen feeders), herbivores (flower sap/tissue feeders), predators (preying or scouting), and combinations of these functions. Different letters indicate significant differences between categories (p < 0.05; see Supplementary Tables S7 and S8).

### Climate and shade-tree canopy drive flower visitor interactions and fruit set

A cocoa pollination process starts from flowering, ends in fruit set, and is mediated by complex interactions between climate, shade-tree canopy cover, and flower visitor community interactions determining cocoa yields (Toledo-Hernández et al., 2017). We used structural equation modelling (hereafter SEM; Lefcheck et al., 2016) to quantify the relationships between (i) flowers per tree, (ii) flower visits (i.e., number of visits by the most frequent visitors) per monitored flower, (iii) fruit set (binary) per monitored flower, (iv) temperature (i.e., mean daily temperature in °C per plot) and (v) shade tree canopy cover (i.e. % canopy cover above cocoa trees per plot). We did not include fertilizer and pesticide application, because these treatments were standardized across all sites and plots in China and application was minimal in Brazil.

We built one SEM for the monitored cocoa production system in China and one for the system in Brazil based on literature-founded hypotheses (Fig. 3a,b, Supplementary Table S9,S10, see “Statistical Analysis” section). In China, increased canopy cover decreases aphid populations in flowers (p<0.001). Fruit set was exclusively driven by increased midge visitation (p=0.007), concurring with previous observation in the cocoa pollination literature (Toledo-Hernández et al., 2017). Furthermore, aphid (p=0.003) and ant (p=0.004) visits were suppressed by the presence of midges in flowers (Fig 3a, Supplementary materials Table 9), linking existing work of ecological interactions between ants and cocoa yields in Asia to pollination (Wanger et al. 2010; Wielgoss et al. 2013). In Brazil, high temperature reduced tree flowering (p=0.005). Additionally, an increased canopy cover (p=0.005) and flowering (p=0.032) led to reduced fruit set, likely due to tree-light competition (Mensah et al., 2023) and energy resources allocated to flowering over fruit set (Valle et al., 1990). The link between increased flowering with thrip visitation (p<0.001) may be a plant-thrip mutualism whereby thrip presence repels flower herbivores in exchange of feeding resources (Shimoda et al., 1997). Unlike in China, increasing ant visitation increased fruit set (p<0.001) and decreased thrip visits in flowers (p=0.001; Fig 3b; Supplementary Table 10).

**Fig. 3.**
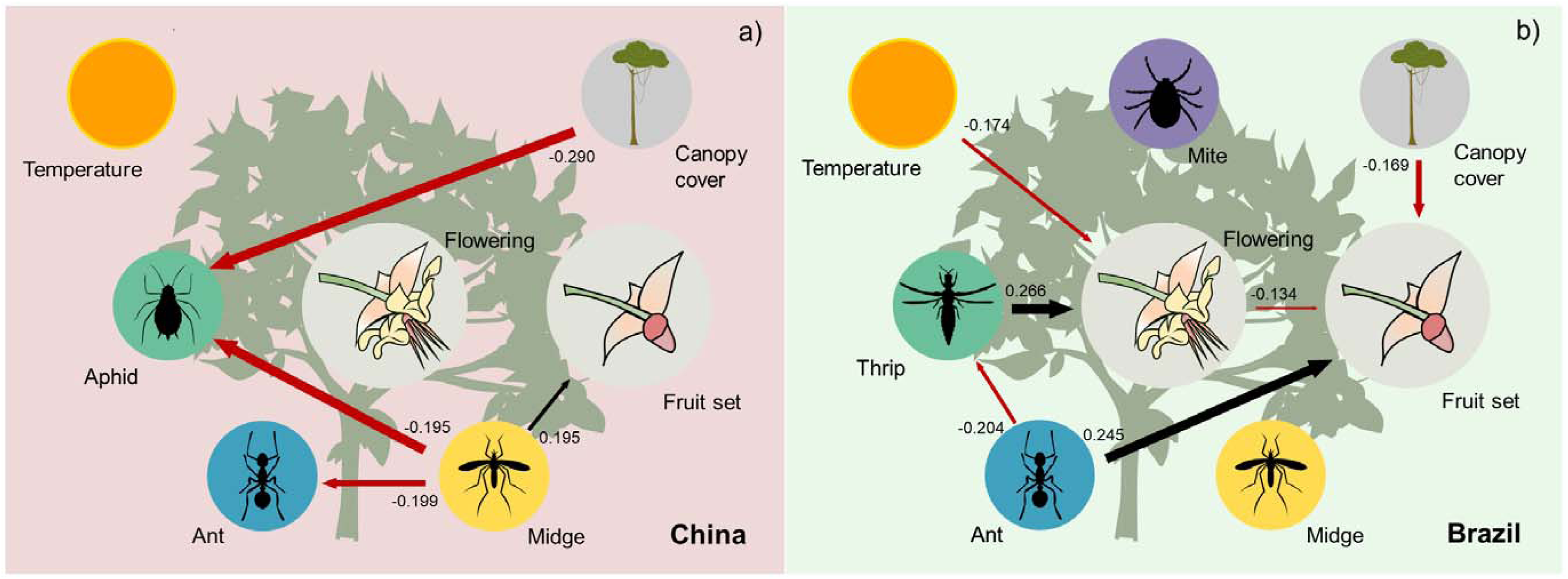
Structural Equation Models (SEMs) on the effects of temperature, canopy cover, and main cocoa flower visitors’ interactions on flowering, and fruit set in China (a) and Brazil (b) cocoa frontiers. Black and red arrows indicate significant positive and negative effects (p < 0.05), respectively (Supplementary materials Table S9, S10). Arrow width is proportional to the strength of the effect, with wider arrows representing lower p-values. Standard coefficient estimates are included.

### Cocoa pollination process – closing the knowledge gap

Our findings indicate that pollination success, and ultimately fruit set driving cocoa yields, is shaped by complex plant-arthropod interactions that are highly variable between continents and different agroforestry systems. In China, we identified three midge morpho-species as main cocoa pollinators (carrying pollen in 29.2% of their visits) and key drivers of fruit set, which is consistent with historical evidence (Wood & Lass, 2008; Toledo-Hernández et al., 2017; Claus et al., 2018). In contrast, it is likely that the scarcity of midges in Brazil led to ant-dominated flower communities, where ants acted as facultative pollinators, pest-predators, and in some cases, flower herbivores. While ant pollination is rare (Beattie, 2006), the high abundance of four morpho-species on cocoa flowers, along with their ability to navigate complex floral structures and carrying pollen in 2% of visits, suggest at least a role in cocoa pollination. Future studies should further investigate the pollination mechanisms of these ants and specifically how their secreted compounds affect pollen viability (Dutton & Frederickson, 2012), as well as their pollination efficacy in self-compatible and self-incompatible trees (Lanaud et al., 2017).

In China, and most cocoa growing regions (Toledo-Hernández et al., 2021; Ocampo-Ariza et al., 2025), aphids are a common pest in cocoa. They proliferated with reduction of shade-tree canopy cover as found in the Americas, Sout East Asia and West Africa (Avelino et al., 2012; Bos et al., 2007; Bisseleua et al., 2013). In both countries, we found that integrating multiple shade tree species benefits nectar and pollen feeding arthropods as potential pollinators, because of enhanced habitat suitability (Toledo-Hernández et al., 2017). These findings on shade-tree diversity and canopy cover enhancing forages in both countries, and reducing herbivores in China, but driving fruit set losses in Brazil, provide initial mechanistic insights into the trade-offs between shade and yields in cocoa production (Tscharntke et al., 2011). However, further research with computer vision approaches needs to elucidate the drivers behind the large variability of shade-tree effects on pollination and pest-control services in agroforestry systems globally (Neither et al., 2020; Lander et al., 2025; Darras et al., 2024).

Flower reductions driven by higher temperature in Brazil highlight the climate risk to cocoa pollination and generally to global cocoa production (Weston, 2025). Indeed, rising temperature, compounded by pollination deficits, contributes to 31% yield losses in Brazil, Ghana, and Indonesia (Lander et al., 2025). Temperature increase could affect pollination through flowering as temperature fluctuations can reduce flower fertility (Raju et al., 2024), delay flowering time (Tun et al., 2021), or shorten its duration (Sheldon, 2019), and thereby create a temporal mismatch with pollinators (Forrest, 2015). Additionally, abnormal temperatures may facilitate the colonization of minor cocoa pests (Oyekodun et al., 2022), such as thrips (Lima et al., 2016), as observed in Brazil. In times where cocoa deficits associated with climate change are jeopardizing the sector globally (Kongor et al., 2024), future studies need to expand work on temperature effects on plant-arthropod interactions across cocoa-growing regions and in sustainable agroforestry systems.

### Supporting pollination and pest-control through climate-smart agroforests

The adoption of climate-resilient agroforestry systems is a promising strategy to future-proof cocoa against rising temperatures (Bunn et al., 2018), and their impact on pollination, biological pest control services, and yield (Lander et al., 2025). Shade-tree canopy management in agroforests helps mitigate temperature fluctuations by stabilizing farm microclimate (Kohl et al. 2024), creating favorable conditions for flowering (Omolaja et al., 2009), flower visitors (Vansynghel et al., 2020), and herbivore natural enemies (Stenchly et al., 2012). Additionally, decaying leaf-litter from shade trees provides organic substrates that enhance flower visitor habitats (Vanhove et al., 2020; Jaramillo et al., 2024). Preserving natural forest and maintaining diversified agroecosystems adjacent to cocoa agroforests facilitates pollinator movements across landscapes (Toledo-Hernández et al., 2021), enhancing their pollination service and boosting farm productivity (Aráujo et al., 2022).

It has, however, also been shown that sustainable cocoa production is not a simple ‘one size fits all’ solution but rather one that requires targeted and location-specific approaches to use pollination for safeguarding productivity under climate change. For instance, more organic matter may also enhance pest and disease pressure in cocoa systems (Wielgoss et al. 2013) and pests and their natural enemies may not always benefit from semi natural habitat in the landscape (Tschantke et al. 2016). Lastly, agroforestry systems may not necessarily help to reduce deforestation, as shown in Southeast Asia (Teo et al. 2025). Thus, more research like the one presented here is crucial to be implemented across all cocoa growing regions to identify locally effective solutions with emerging technologies. Only then will we be able to safeguard agrobiodiversity and productivity in sustainable systems under climate change (Wanger et al. 2020).

## Materials and methods

### Study area

We conducted our study in emerging and consolidated cocoa frontiers, respectively China and Brazil, between March and July (China) and between May and August (Brazil) 2023. In China, cocoa cultivation was first introduced in the 1960s to the Hainan region, the major national supplier of cocoa, where production remains at a very small scale (Zhao et al.,2018). While total cultivated area reached only 600 ha in 2018, national experts forecast a rapid expansion in unproductive agricultural land to meet the increasing national chocolate demand (Li et al. 2022). Our research was conducted in the Xinglong Tropical Botanical Garden (18°44’01” N, 110°11’38” E) in Hainan Province (Supplementary Figure S4a). The region is characterized by a tropical monsoon climate with a mean annual temperature and precipitation of about 25°C and 2,103 mm, respectively (HPLHO, 2004). While most cocoa is cultivated under full sun, agroforests with coconut (*Cocos nucifera* L.) and areca (*Areca catechu* L.) palms are expanding and are intercropped with high-yielding cocoa varieties (Zhang. 2021).

In contrast, Brazil is a historical exporter of cocoa with production dating back to the 18^th^ century (Rolim and Chiarello, 2004; Ruf and Schroth, 2004). In 1989, the country experienced a dramatic production bust (Clough et al., 2009) associated to the witches’ broom (*Moniliophthora pernicosa* L.) disease outbreak (Cassano et al., 2009) that led to an unprecedented economic crisis lasting until today (Marques et al., 2021). We conducted our study in the municipality of Ilhéus (14° 40’ 16.08’’ S, 39° 12’ 14.39’’ W) in Southern Bahia (Supplementary Figure S4b). This region is characterized by a humid tropical climate without a defined dry season and mean annual temperature of 25°C and precipitation of 1,500 mm (Heming et al., 2022). Here, cocoa grows primarily under the shade of native trees from the Atlantic Forest, an agroforestry system known as cabruca, highly important for its biodiversity conservation value (Cassano et al., 2012). Climate change and socio-economic pressures are pushing conversion of cabrucas into full sun cocoa, or other intensified land uses, jeopardizing the regional ecological and economic stability (Gateau-Rey et al., 2018).

### Study design, shade-tree diversity and canopy cover

Our study aimed to investigate how climate (mean daily temperature), shade-tree diversity and canopy management within the plot influences flower visitors, their ecological functions, and the resulting effects on fruit set as a precursor of cocoa yield.

In China, we conducted research on three cocoa farms representative of the study area and each reflecting a different level of shade-tree diversity and canopy management: (i) a monoculture (unshaded system), (ii) a low-shade system (shaded with areca palms and <30% canopy cover area), and (iii) an intermediate-shade system (shaded with coconut palms and 40–50% canopy cover area; Supplementary Table S1). The farms had an area < 2 ha and were at a minimum distance of 100 m between each other. They were also treated at the same time and with the same quantities of fertilizers and pesticides and pruned uniformly by the same trained support staff. In each farm, we established three (10 x 10 m) plots at a minimum distance of 10 m between plots, resulting in a total of nine plots for the study. (Supplementary Figure S4c).

In Brazil, we selected 12 cabruca plots of 40 x 40 m at least 500 m apart from each other and within cocoa farms ranging from 5-202 ha. Besides the smaller production area (5 ha) owned by one smallholder, all cabrucas were managed by cooperatives, with one person responsible for each plot. All plots were rainfed, cocoa trees pruned, and with limited to no use of fertilizer. Within our 12 plots, we differentiated between three cabruca types based on their shade-tree conservation status: i) low-shade cabrucas with shade trees removal (1-3 shade tree species and 30-40% canopy cover area), ii) intermediate-shade cabrucas, with moderate shade tree removal (4-5 shade tree species and 40-50% canopy cover area), and iii) high-shade cabrucas with minimum to no shade tree removal (7-9 shade tree species and >50% canopy cover area; Supplementary Table S1; Figure S4d).

On all 21 plots, we recorded species identity and densities of all shade trees. We then recorded shade tree canopy cover as mean percentage using five photographs randomly taken within the plots at 5 m height and processed them in Image-J software (Wanger et al., 2010).

### Embedded computer vision cameras

We used portable embedded computer vision (ECV) cameras (Darras et al., 2024) to achieve continuous flower visitor monitoring over an average operation time of 19.1 hours, covering nearly half the time in which the flower is receptive to pollination (Toledo-Hernández et al., 2017).

Our ECV camera uses an openMV H7+ microcontroller board coupled to a CMOS image sensor board, together with a power management and control system (Balle et al. 2024) encapsulated in a waterproof housing. We mounted a M12-mount lens with 8 mm focal length and F2.0 aperture in front of the 5-megapixel image camera sensor, to achieve macro images of even the smallest (< 3 mm; Wolcott et al., 2023) cocoa flower visitors. Since flower visitors could occur during the day and at night, we used the camera’s built-in near-infrared LEDs to illuminate our close-range target at night. We removed the infrared cut-off filter from the back of our lenses to be able to utilize that illumination at night. All images were captured as near-infrared photographs but saved in grayscale color (256 levels). The camera’s three internal rechargeable lithium batteries (18650 size) provided a total capacity of 10 Ah, powering the system for up to 24 hours in continuous monitoring mode. In China, where all components were easily available, we extended the runtime of the device to maximize flower visitor monitoring duration of up to 48 hours with an external 13.4 Ah power bank connected to the camera’s USB type-C port (Supplementary Figure S5a,b).

The ECV camera runs a continuous loop of capturing an image, applying frame differencing, and executing a “blob” detection algorithm: This algorithm detects blobs as areas with light and/or shade changes in the current image by comparing it to a reference image (Lindenberg, 1998; Manchanda and Shanu, 2016). We used Lab color space thresholds of [(0,3), (−3,3), (−3,3)] and a minimum blob size of 11,250 pixels to determine whether a blob triggers a detection. To avoid false detections from changing light conditions or growing flowers, the reference image is updated every 60 seconds by blending it with an alpha value of 50% with a new frame if there are currently no detections. This computer vision method is suitable for detecting moving targets, like cocoa flower visitors, and produces high precision results on static and monotone image backgrounds with precisely adjusted area and color thresholds (Darras et al., 2024). In our case, however, the fragile flowers would often move due to light breezes, causing the camera to trigger a false detection (∼95% images with no flower visitor) as the flower position would change and light and shade patches in the image would also move.

When one or more moving targets, such as flower visitors, are detected in the captured image, the image is saved as a grayscale JPEG onto the flash memory card. The image data, such as the date, time, exposure, gain, and frame rate are added in a CSV file. Concurrently, all detected blobs in the image are saved, with their size, position, and color range documented in another CSV. A status information line is added to a CSV file every 10 minutes to monitor the camera operation, including details about the battery voltage, charging status, core temperature, and the reference voltage of the processor chip (Supplementary Figure S5c). These output files, in combination of manual annotations of flower visitor presence in the stored images, allow for extraction of monitoring duration, total flower visitor detections, and time spent in flowers at the end of a deployment.

### Camera trap deployment, and fruit set recording

First, we selected and marked nine and six cocoa trees per plot in China and Brazil, respectively. Selected trees were of similar size and age, and in general not affected by pests and diseases. We used the marked trees for the camera deployment, which involved three steps: i) mounting the camera to a marked tree branch using a flexible-arms tripod, ii) focusing the ECV camera on a newly opened flower at a distance of <3 cm and with a frontal view on the flower; and iii) setting up the blob detection algorithm in a continuous (day and night) monitoring mode. In each monitoring day, we deployed a maximum of nine cameras in China and 12 cameras in Brazil, distributed across the marked trees and plots. As the number of flowers available was uneven across the trees and plots, our dataset is unbalanced for the number of flowers monitored within trees and plots. After the camera installation, we labelled the monitored flowers with waterproof adhesive stickers pinned near the flower and counted the number of open flowers in each monitored tree. We retrieved the device once its battery was depleted (as indicated by the lack of LED signals), and enclosed flowers with small mesh bags to avoid further visitation. We then recorded the fruit set of the monitored flowers when they did not abscise after approximately 48 hours (Almeida et al., 2007) and showed bud formation in the following seven days.

### Flower visitor identification and ecological function classification

We manually sorted the images stored in the camera’s memory card. Our image post-processing approach consisted of five steps. First, we sorted images with flower visitors and without flower visitors (Supplementary Figure S6a), giving a ratio of ∼5:95. Second, we used the true positive images and identified visitors to order, family and – where possible, morpho-species level (Supplementary Figure S6b). Third, when possible we classified visitor images into three functional groups (i.e. nectar and pollen foragers, flower-herbivores, arthropod-predators) and their combinations (i.e. foragers/herbivores, foragers/predators, herbivore/predators) following their visitor behavior as observed in the saved images sequence, and recorded if cocoa pollen was present on the flower visitor body (Supplementary Figure S3, S6c, Table S5). We classified visitors as foragers when they fed on nectar or pollen and visited female and male reproductive sections of the flower. For herbivores, visitors were piercing and cutting floral tissues while predators were actively hunting or scouting (i.e. moving across the flower and pausing intermittently) for prey. Many morpho-species showed combined behaviors like nectar and pollen as well as plant sap/tissue feeding (i.e. forager/herbivore), foraging and predation (i.e. forager/predator), and herbivory and predation (i.e. herbivore/predator). Fourth, we quantified number of visits, counting a new visit every time a visitor appeared touching the flower petals, staminodes, style or anthers with a time difference of at least 30 seconds from the previous flower visit of the same species/morpho species to minimize the chance of pseudo replication (Supplementary Figure S6d). Number of visits should be interpreted with caution, since some visitors move rapidly and the low camera framerates may have missed some visits.

Finally, we calculated the duration of flower visits using the first and last detection time-data from the metadata of the stored images (Supplementary Figure S6e).

### Climate data

We used the ERA5-Land reanalysis dataset from 1950 to the present (Muñoz Sabater, 2019), with a spatial resolution of 11 km, to extract the daily mean air temperature at 2 m above the land surface for our 21 plots in China and Brazil. Extraction was performed for the deployment period at the plot level (), using the coordinates of each plot. Plots located less than 11 km apart were assigned to the same temperature value, and temperature values, originally provided in Kelvin (K), were converted to Celsius (°C). To corroborate that the ERA5-Land dataset accurately reflects the climate conditions at our plots, we compared it with a subset of temperature measurements recorded by dataloggers placed in the center of the nine plots in China where devices were available (R=0.76, p<0.001; Supplementary Figure S7).

### Statistical analysis

We carried out all statistical analysis in R development software (R Core Team, 2024). First, we investigated the effect of increasing shade-tree species richness and canopy cover across different agroforestry systems (i.e. monoculture, low, moderate shade systems in China; and low, intermediate, and high shade cabrucas in Brazil) on flower visitor functional groups (i.e. proportion of visits by foragers, herbivores, predators, and their three combinations). Here, we used the proportion of visits of each group aggregated at the country, agroforestry system, plot and tree level. Then, we used the glmmTMB package (Brooks et al., 2023) to build three generalized linear mixed effect models (hereafter “GLMMs”) for each functional group as response, and agroforestry system as predictor variables. We fitted our models with a negative binomial distribution to account for overdispersion, and separately country and plot as random effects. We then carried out a multiple comparison of means analysis using the functions pairwise() and adjusted tukey() from the “emmeans” (Lenth, 2022) and “multcomp” (Hothorn et al., 2008) R-packages.

Second, we used Structural Equation Models (SEMs) to investigate the direct effects between temperature (i.e. mean daily temperature), canopy (i.e. percentage of shade tree canopy cover) on flower visitors’ interactions (i.e. proportion of visits by the main flower visitor taxonomic groups) and its relation to flowering (i.e. number of tree flowers), and fruit set (i.e. 1 = yes, 0 = no). All our interacting variables were sorted by country, agroforestry system, plot, tree, and monitoring day, except temperature which was measured at the plot and monitoring day, and canopy cover at the plot level. We used the piecewiseSEM package (Lefcheck, 2016) to build six individual linear equations for each country to evaluate these interactions. We fitted linear models for our SEMs to reduce complexity and avoid convergence issues (Pearl, 2016). For China, we included the following models which reflect our hypotheses:

(a) lm (flowering ∼ temperature + canopy + aphid + ant + midge)
(b) lm (fruit set ∼ temperature + canopy + flowering + aphid + ant + midge)
(c) lm (aphid ∼ temperature + canopy + ant + midge)
(d) lm (ant ∼ temperature + canopy + midge)
(e) lm (midge ∼ temperature + canopy)

For Brazil, the models were:

(f) lm (flowering ∼ temperature + canopy + thrip + ant + mite + midge)
(g) lm (fruit set ∼ temperature + canopy + flowering + thrip + ant + mite + midge)
(h) lm (thrip ∼ temperature + canopy + ant + mite + midge)
(i) lm (ant ∼ temperature + canopy + mite + midge)
(j) lm (mite ∼ temperature + canopy + midge)
(k) lm (midge ∼ temperature + canopy)

We hypothesized that flowering (models a, f) and fruit set (models b, g) are influenced by increased temperature (Raju et al., 2024; Lander et al., 2025), canopy cover (Adjaloo et al., 2012; Niether et al., 2020), and the ecological functions of the main flower visitors (Bronstein et al., 2007). Flower visitors may act as pollinators (e.g., midges), herbivores and predators (e.g., mites and ants), or perform multiple roles (e.g., foraging and herbivory by aphids and thrips), thereby affecting flowering and fruit set through their use of floral resources. Flowering may also influence fruit set by increasing resource availability for pollinators (Ulyshen et al., 2023) and/or via selective abortion of flowers and early fruits (Sutherland, 1986; Agustí & Primo-Millo, 2020). Finally, the visitation number of each visitor group (models c-e, h-k) could be shaped by temperature (Halsch et al., 2021), canopy cover (Vansynghel et al., 2022), and interactions among visitors according to their ecological functions. For instance, ants may prey on arthropods and/or facilitate potential pests (Klein et al., 2002) such as aphids and thrips, which can in turn affect pollinator visitation (LeVan & Holway, 2015) by midges. Most of these relationships have not yet been disentangled for cocoa.

Prior to model fitting, we evaluated the normality assumption of the residuals for our response variables using the Shapiro-Wilk normality test. All variables were transformed using the “bestNormalize” work-package, which selects the optimal transformation for each variable. (Peterson, 2021). Models for both countries were saturated, including all plausible causal links among the factors. Because saturated models leave no conditional independence, Fisher’s C cannot be computed, and global goodness-of-fit is undefined. We chose this approach to capture all potential interactions in our system, which are not fully understood, and interpret results based on the estimated path coefficients (Supplementary Table S11, S12). Additionally, following the same model structure as above, we built GLMM models with negative binomial distribution, except for fruit set which was treated binomial family, and plot as random effect, to validate our linear model results (Supplementary Table S11, S12).

---

**Figure S1.**
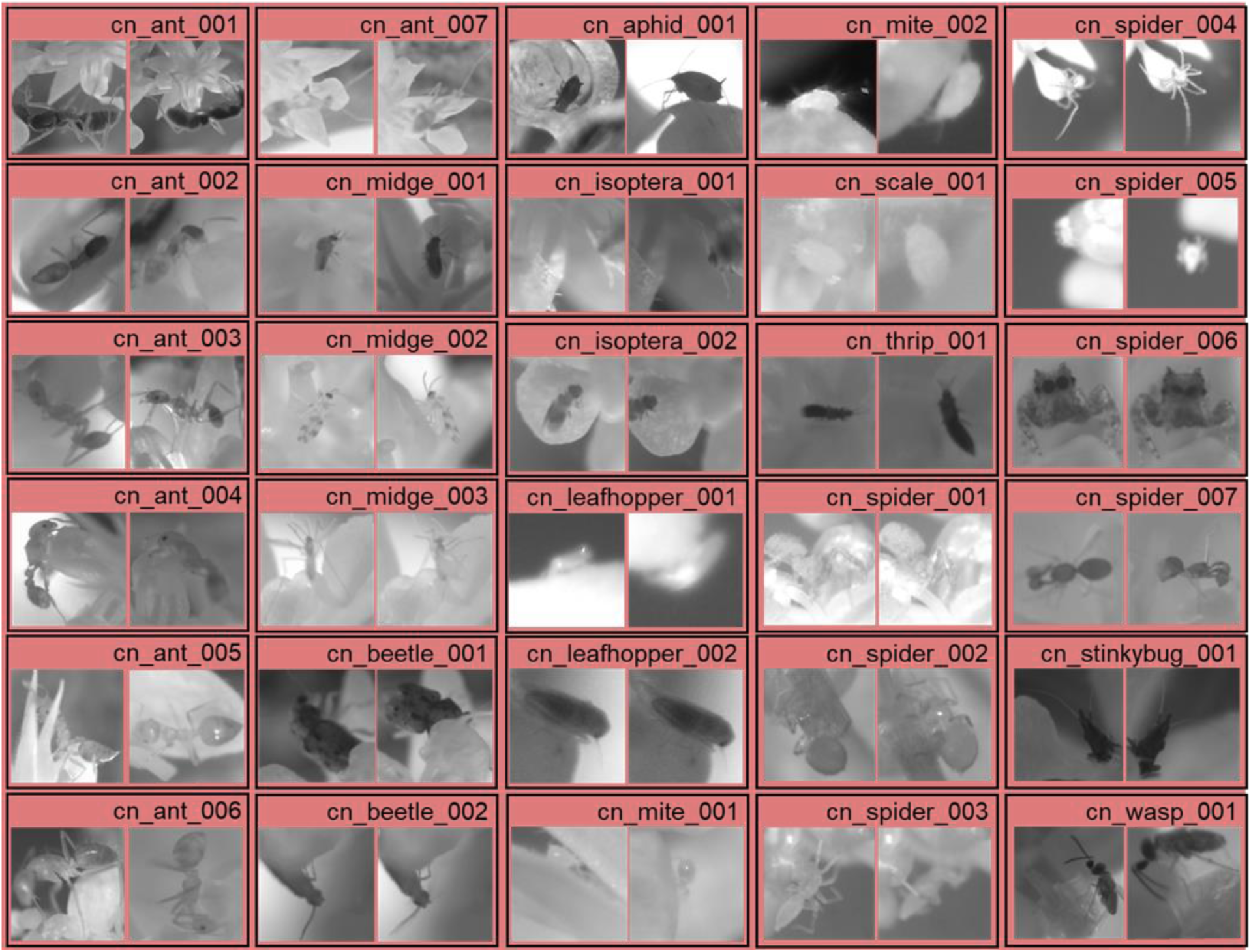
Flower visitor morphospecies of cocoa systems in Hainan, China. We identified 30 morphospecies in 12 taxonomic groups. Midge morphospecies (cn_midge_001-003) carried pollen in 29:2% of the visits, resulting in key drivers of fruit set.

**Figure S2.**
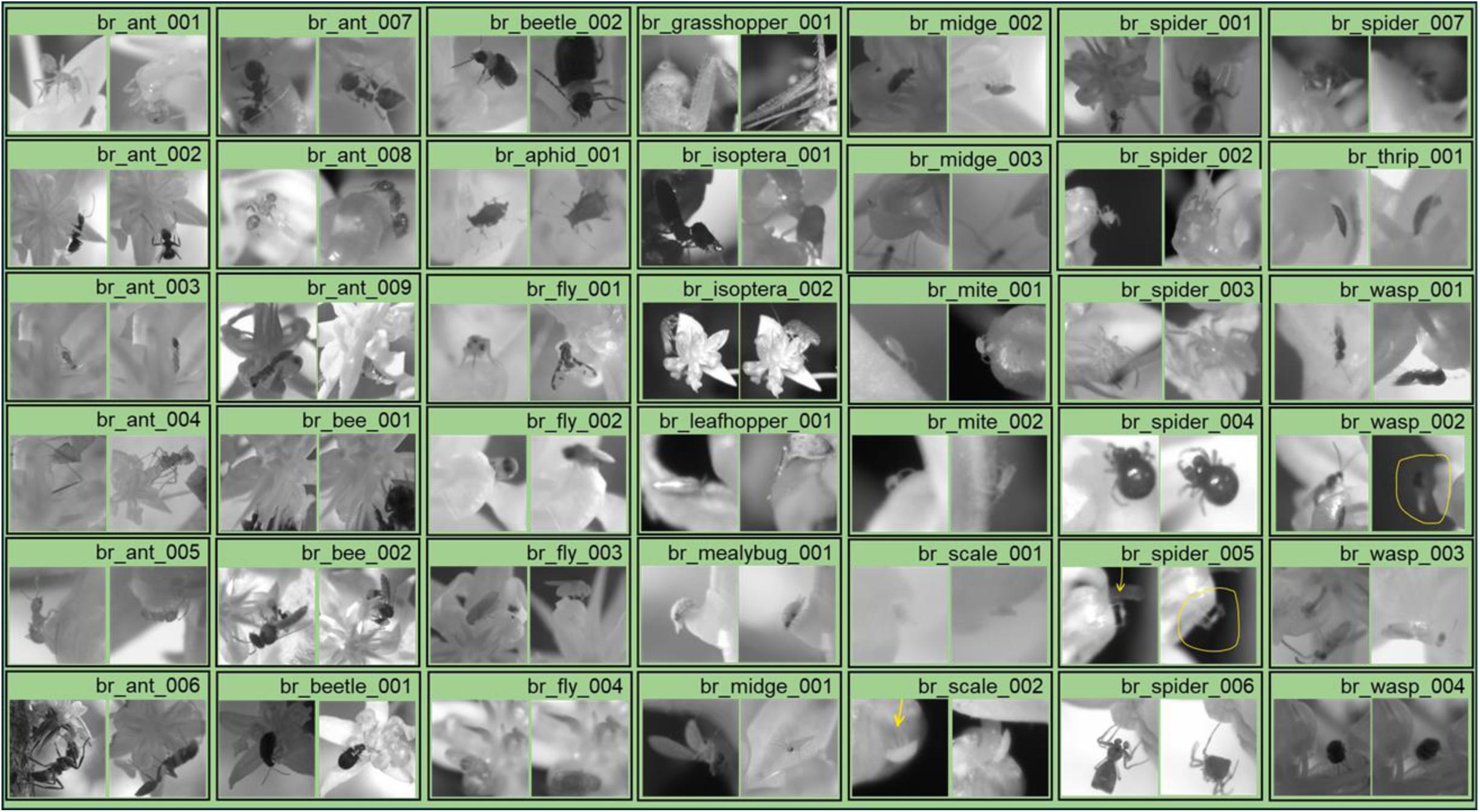
Flower visitor morphospecies of cocoa systems in Bahia, Brazil. We identified 42 morphospecies in 15 taxonomic groups. Ant morphospecies (br_ant_003,005,007,008) carried pollen in 2% of the visits, resulting in key drivers of fruit set.

**Figure S3.**
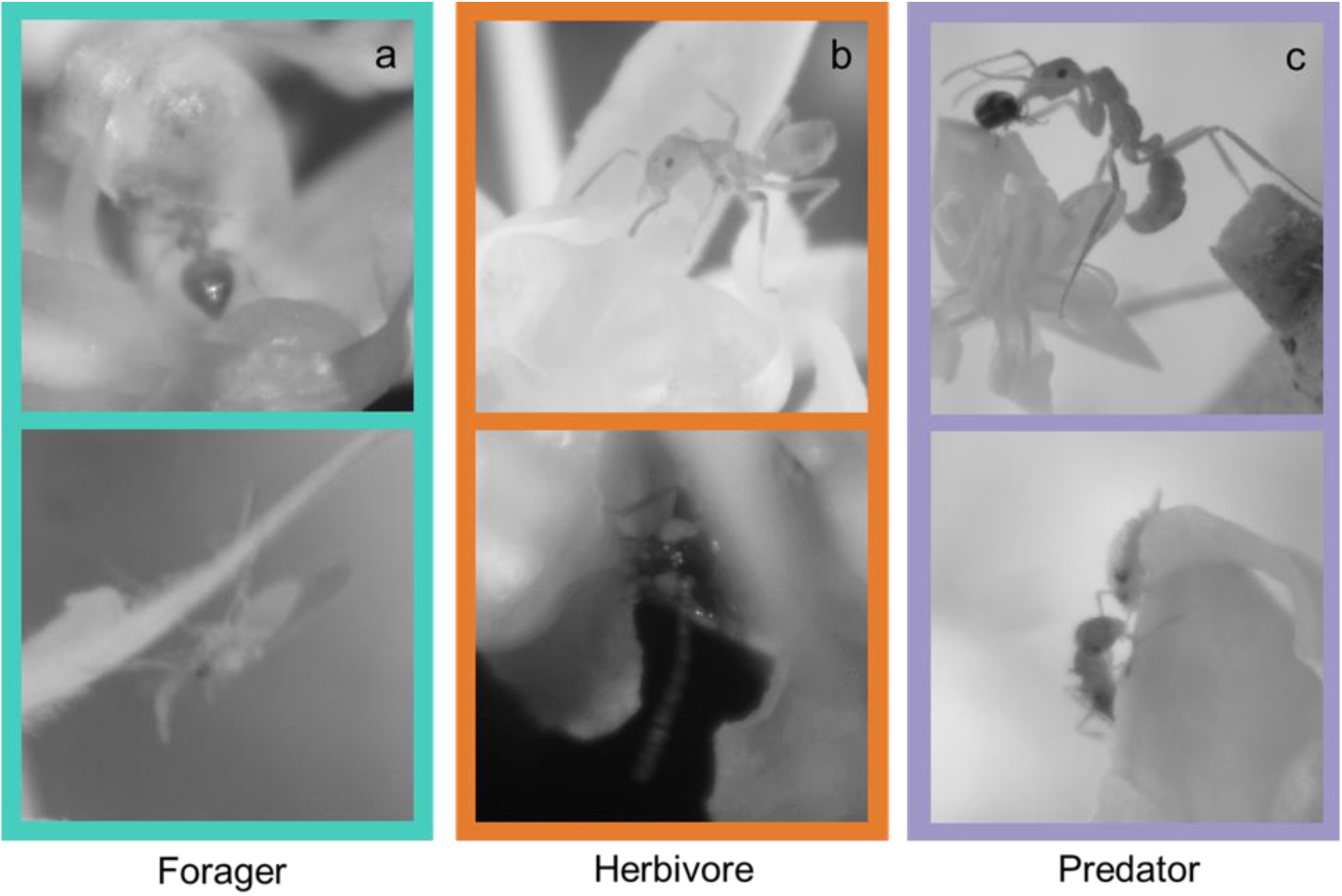
We recorded three basic ecological functions among flower visitors. Namely, (i) nectar and pollen foraging, (ii) flower sap and tissue herbivory, and (iii) predation; and three combined functions (iv) forager/herbivore, (v) forager/predator, and (vi) herbivore/predator. The images illustrate examples of the basic functions: an ant feeding on pollen from an anther and a midge feeding on nectar on a staminode (forager, a); an ant cutting and a beetle feeding on flower petals (herbivore, b); and an ant attacking a beetle and chasing a mealybug (predator, c). Some flower visitors performed combined functions: iv) forager/herbivore, v) forager/predator, vi) herbivore/predator. A detailed explanation of the classification protocol is provided in Supplementary Table S5.

**Figure S4.**
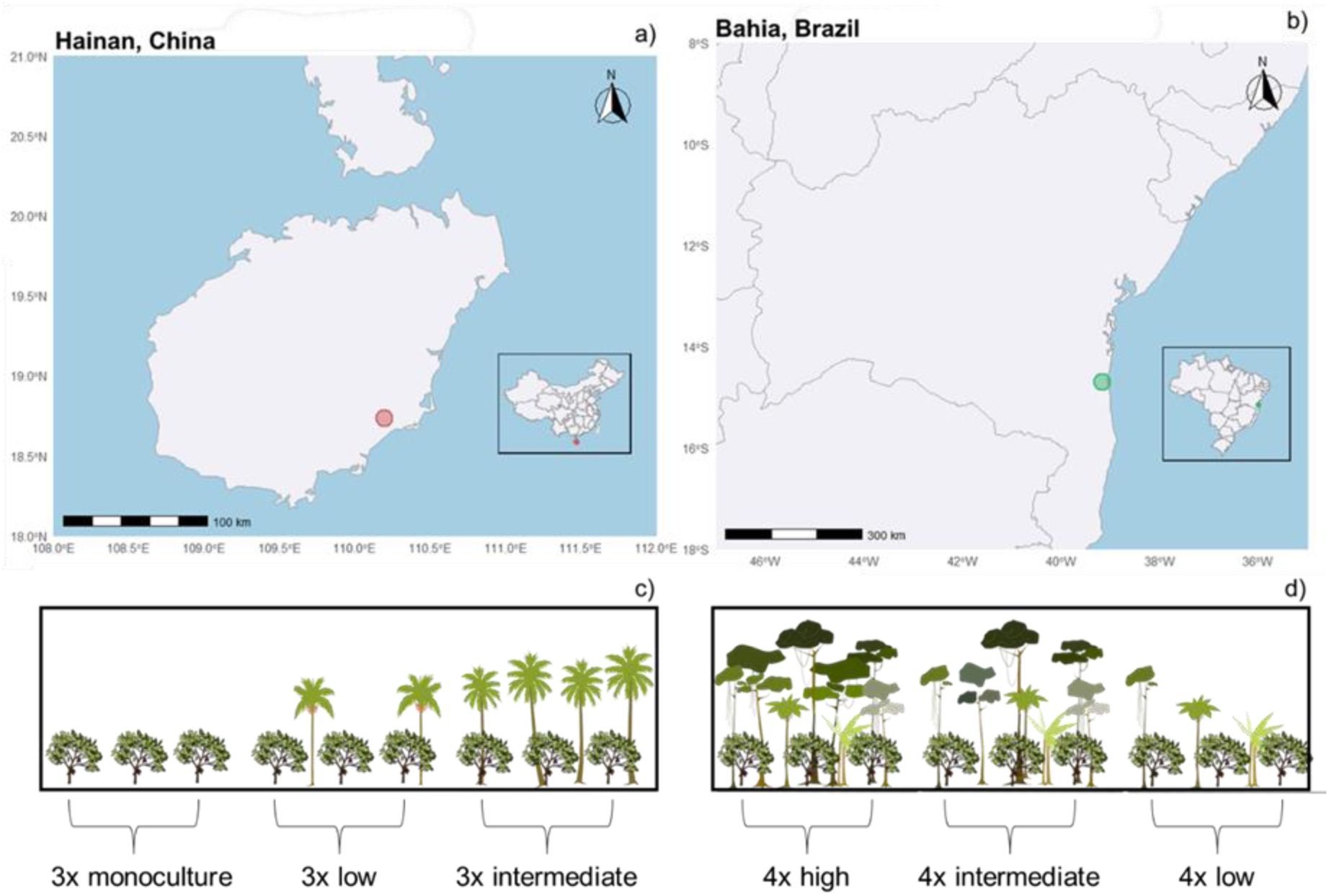
We conducted the monitoring of cocoa flower visitors in two regions: Hainan, China (red dot; a), and Bahia, Brazil (green dot, b). In each region, we selected cocoa farms representing contrasting shade-tree diversity and canopy cover management. In Hainan the three farm types were: (i) Monoculture, with no shade (unshaded system); (ii) Low-shade system, shaded by areca palms with less than 30% canopy cover; and (iii) Intermediate-shade system, shaded by coconut palms with 30–40% canopy cover (c). In Bahia the three farm types were: (i) Low-shade cabrucas, which had undergone heavy shade tree removal (1-3 shade tree species and between 30-40% canopy cover); (ii) Intermediate-shade cabrucas, with moderate shade tree removal, or 4–5 shade tree species and 40-50% canopy cover); and (iii) High-shade cabrucas, with minimal or no shade tree removal, or 7-9 shade tree species and over 50% canopy cover (d). We established three replicates per system type in China and four replicates per system type in Brazil.

**Figure S5.**
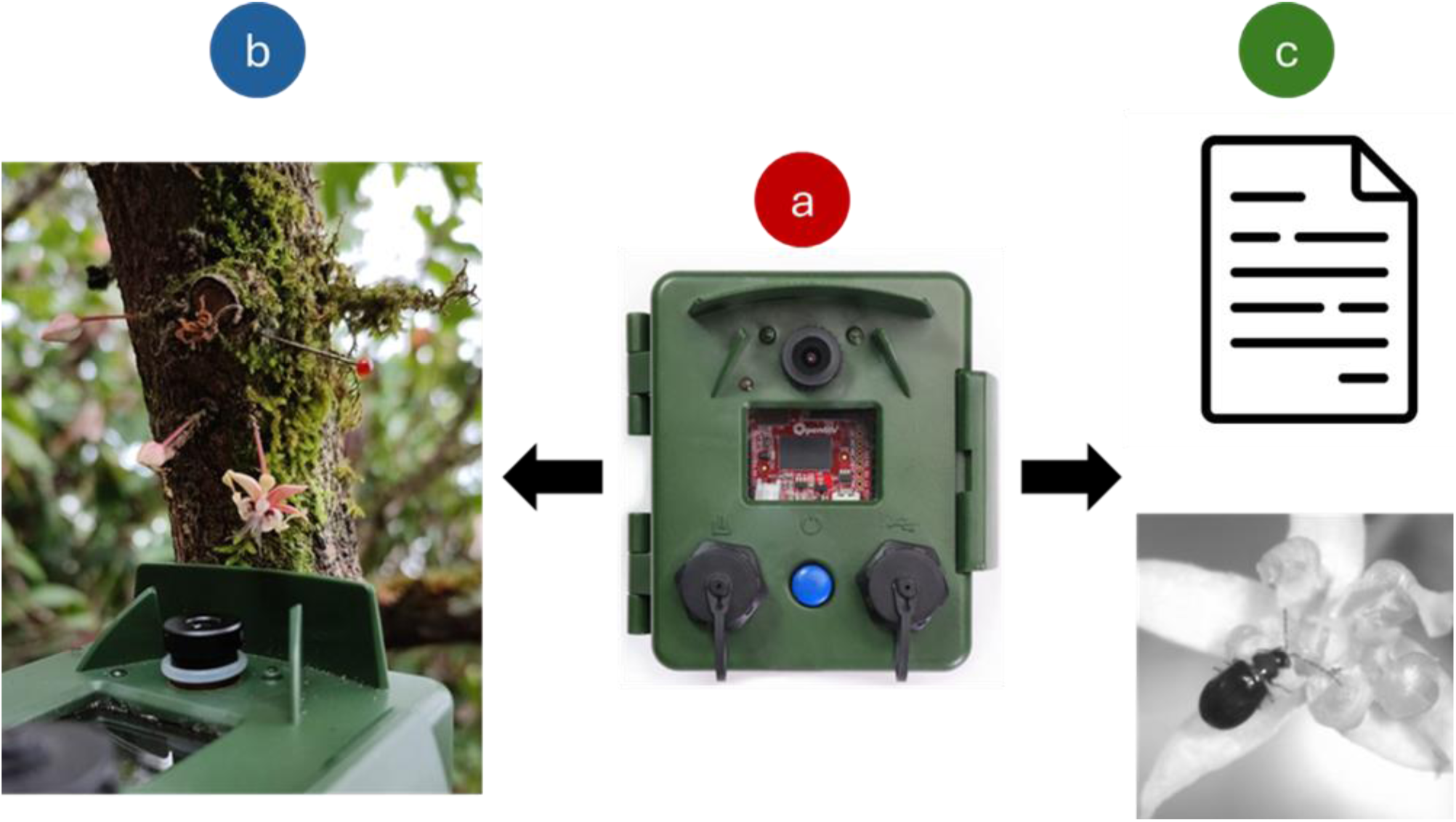
Our embedded computer vision (ECV) camera trap used to monitor cocoa flower visitors in China and Brazil. The ECV system (Balle et al., 2024) is built with OpenMV H7 Plus microcontrollers and an image sensor board housed in a waterproof casing, equipped with an M12 lens (8 mm focal length, F2.0 aperture; a). Cameras were deployed directly on cocoa flowers and left to operate until battery depletion (b). The ECV system applies frame differencing and blob detection algorithms, which are well-suited for detecting moving targets such as cocoa flower visitors. When motion is detected, the system saves an image to the flash memory card, along with a CSV file with the date, time, and camera operation status (c). These files allow us to assess flower monitoring duration, total visitor detections, visitation time, and visitor behavior.

**Figure S6.**
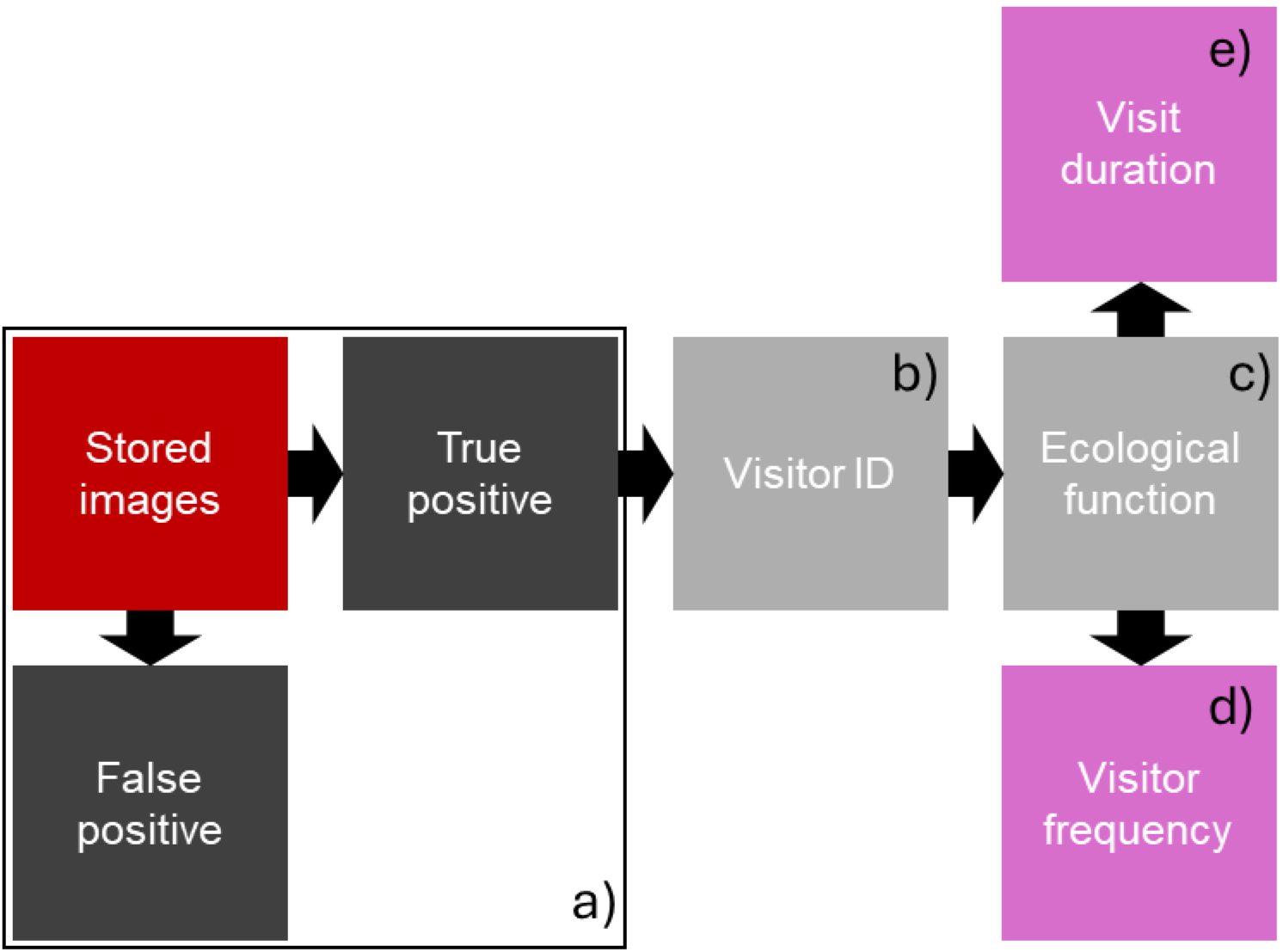
Step-by-step diagram of flower visitor image sorting. First, all images stored on the flash memory card were classified into two categories: *false positives* (images without visitors) and *true positives* (images containing at least one visitor, a). Second, we identified the visitors in true positive images to taxonomic groups (i.e. typically at the order, family, and morphospecies levels when possible, see Figure S1,S2, b). Third, we categorized the identified visitors into three basic ecological functions and their combinations based on their observed behavior (see Supplementary Figure S3, Supplementary Table S5). Fourth, we quantified visits by counting a single visit each time a visitor was observed contacting flower parts (i.e. petals, staminodes, style, or anthers), provided there was a time gap of at least 30 seconds since the previous visit by the same morphospecies (d). Finally, we calculated the duration of each visit using date and time from the CSV file associated with each image (e).

**Figure S7.**
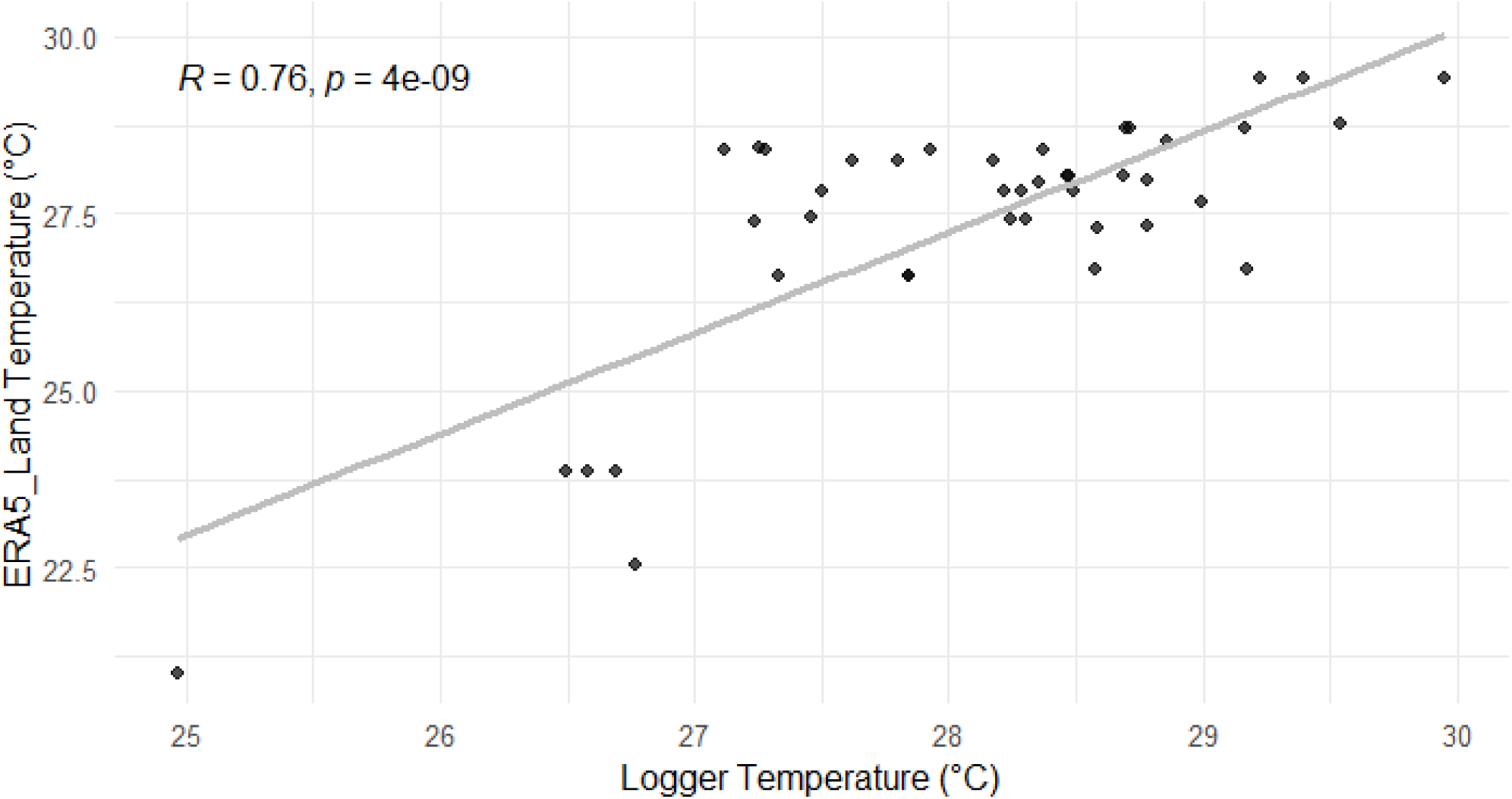
Correlation between Mean daily temperature (°C) from plot-level loggers and spatiotemporal (ERA5_Land) datasets in our nine research plots in China. Loggers recorded temperature every 30 minutes at 1.5 m above ground from May to July 2023, and daily means were calculated and compared with the corresponding spatiotemporal data.

**Table S1.**
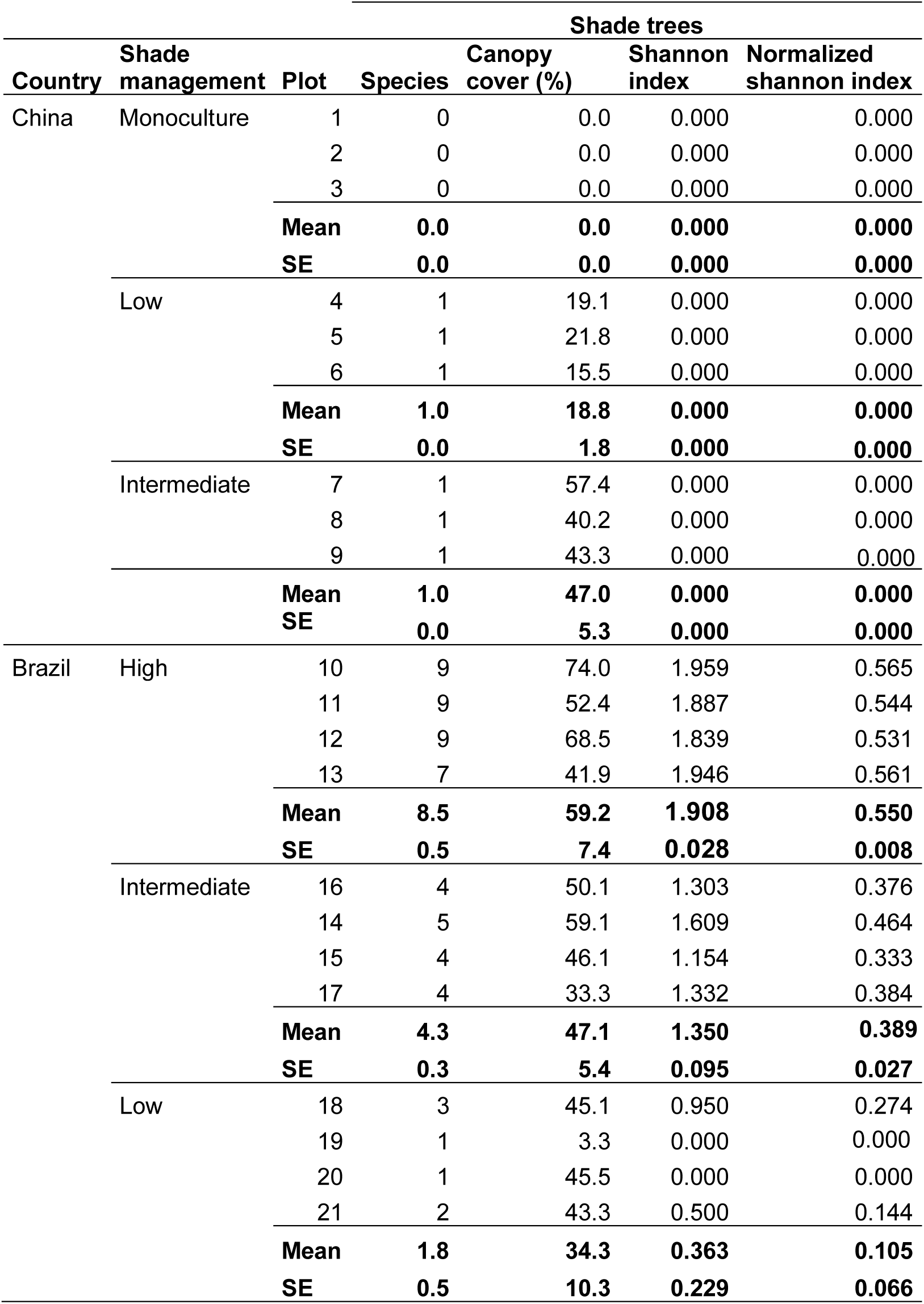
Summary of shade-tree species diversity and canopy cover management in our 21 research plots in China and Brazil.

**Table S2.**
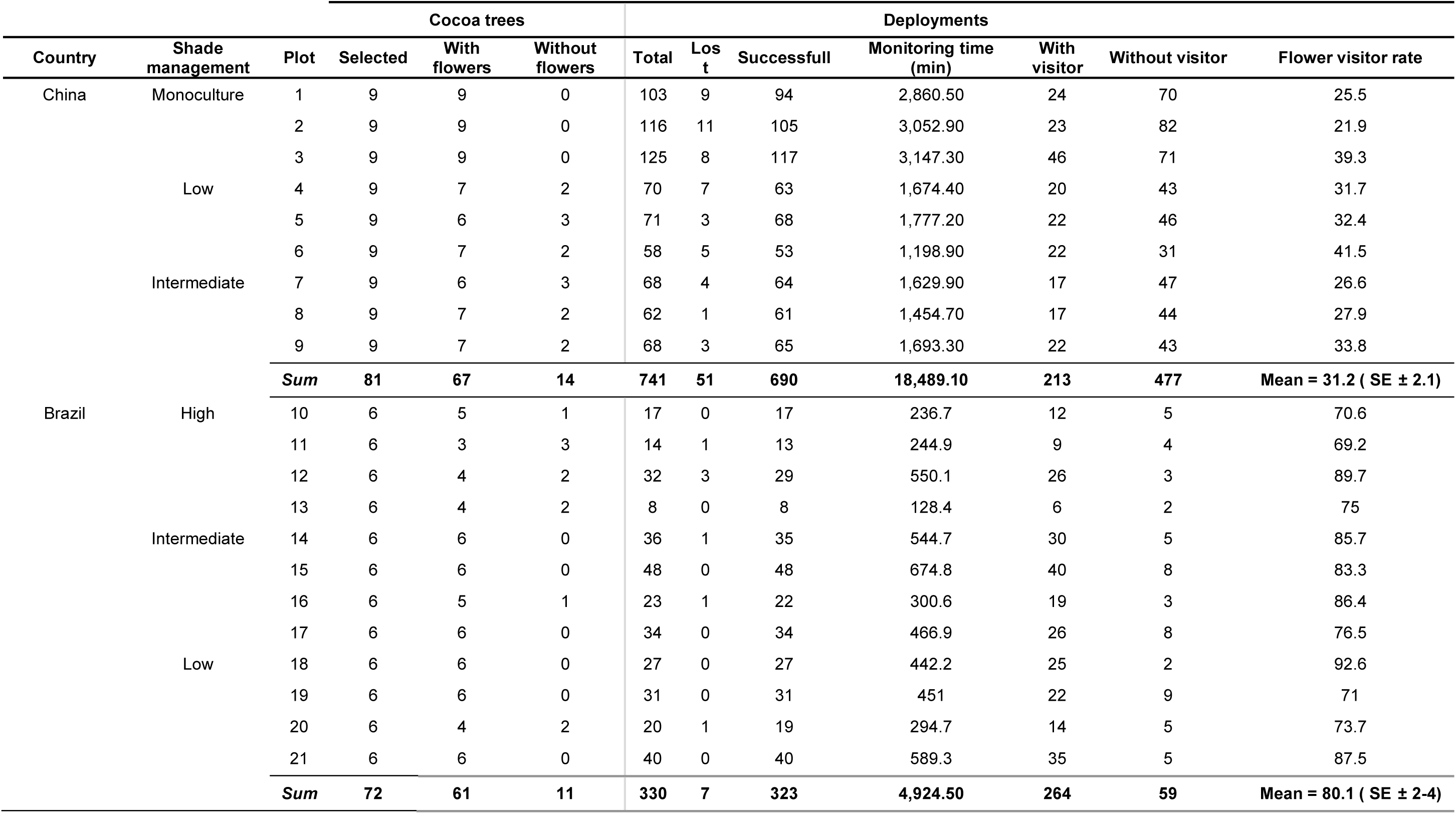
Summary of ECV camera deployments in the 21 research plots in Hainan, China and Bahia, Brazil.

**Table S3.** Summary of visits sorted by taxonomic groups recorded with ECV cameras in our 21research plots in Hainan, China and Bahia, Brazil. Immature stage visitors (i.e. larvae, nymph) and not identified (i.e. NI) visitors are not included in the sum of visits (i.e. Visits).

| Country | Shade management | Visits | Mite | Spider | Isoptera | Midge | Fly | Beetle | Aphid | Leafhopper | Scale | Mealybug | Stinkybug | Bee | Ant | Wasp | Grasshopper | Thrip | Larvae | Nymph |
| --- | --- | --- | --- | --- | --- | --- | --- | --- | --- | --- | --- | --- | --- | --- | --- | --- | --- | --- | --- | --- |
| China | Monoculture | 134 | 35 | 4 | 0 | 4 | 0 | 1 | 12 | 4 | 14 | 0 | 0 | 0 | 59 | 0 | 0 | 1 | 0 | 0 |
|  |  | 145 | 9 | 0 | 0 | 11 | 0 | 0 | 69 | 17 | 0 | 0 | 0 | 0 | 36 | 0 | 0 | 1 | 1 | 1 |
|  |  | 367 | 1 | 0 | 0 | 24 | 0 | 0 | 237 | 2 | 0 | 0 | 0 | 0 | 98 | 2 | 0 | 0 | 3 | 0 |
|  | Low | 54 | 1 | 3 | 0 | 14 | 0 | 0 | 8 | 0 | 2 | 0 | 0 | 0 | 23 | 3 | 0 | 0 | 0 | 0 |
|  |  | 108 | 7 | 2 | 1 | 17 | 0 | 0 | 26 | 0 | 0 | 0 | 0 | 0 | 54 | 1 | 0 | 0 | 0 | 0 |
|  |  | 74 | 0 | 3 | 0 | 23 | 0 | 0 | 17 | 0 | 0 | 0 | 1 | 0 | 23 | 0 | 0 | 0 | 1 | 6 |
|  | Intermediate | 39 | 0 | 8 | 0 | 10 | 0 | 1 | 4 | 2 | 0 | 0 | 0 | 0 | 12 | 2 | 0 | 0 | 0 | 0 |
|  |  | 50 | 6 | 16 | 0 | 14 | 0 | 0 | 1 | 1 | 0 | 0 | 0 | 0 | 12 | 0 | 0 | 0 | 0 | 0 |
|  |  | 59 | 0 | 6 | 1 | 15 | 0 | 0 | 2 | 0 | 0 | 0 | 0 | 0 | 32 | 0 | 0 | 3 | 0 | 0 |
|  | **Total** | 1030 | 59 | 42 | 2 | 132 | 0 | 2 | 376 | 26 | 16 | 0 | 1 | 0 | 349 | 8 | 0 | 5 | 5 | 7 |
|  | **Mean** | 114.4 | 6.6 | 4.7 | 0.2 | 14.7 | 0.0 | 0.2 | 41.8 | 2.9 | 1.8 | 0.0 | 0.1 | 0.0 | 38.8 | 0.9 | 0.0 | 0.6 | 0.6 | 0.8 |
| Brazil | High | 293 | 5 | 13 | 2 | 0 | 1 | 22 | 10 | 0 | 1 | 0 | 0 | 0 | 102 | 1 | 1 | 135 | 0 | 0 |
|  |  | 67 | 5 | 1 | 0 | 1 | 1 | 4 | 0 | 0 | 0 | 0 | 0 | 0 | 19 | 0 | 0 | 36 | 0 | 0 |
|  |  | 164 | 13 | 0 | 0 | 0 | 0 | 2 | 0 | 0 | 0 | 1 | 0 | 0 | 54 | 0 | 0 | 94 | 0 | 0 |
|  |  | 75 | 1 | 0 | 0 | 1 | 0 | 0 | 0 | 0 | 0 | 0 | 0 | 0 | 23 | 0 | 1 | 49 | 0 | 0 |
|  | Intermediate | 202 | 34 | 0 | 0 | 3 | 1 | 3 | 0 | 0 | 0 | 1 | 0 | 0 | 83 | 5 | 0 | 72 | 0 | 0 |
|  |  | 494 | 14 | 1 | 0 | 2 | 2 | 43 | 0 | 0 | 7 | 2 | 0 | 1 | 50 | 0 | 0 | 372 | 0 | 0 |
|  |  | 488 | 85 | 3 | 0 | 0 | 1 | 3 | 7 | 24 | 1 | 6 | 0 | 0 | 193 | 1 | 0 | 163 | 0 | 1 |
|  |  | 352 | 33 | 0 | 0 | 4 | 1 | 6 | 4 | 1 | 2 | 0 | 0 | 0 | 120 | 2 | 0 | 179 | 0 | 0 |
|  | Low | 111 | 2 | 0 | 0 | 0 | 0 | 0 | 0 | 0 | 0 | 0 | 0 | 0 | 65 | 1 | 0 | 32 | 10 | 1 |
|  |  | 220 | 17 | 0 | 0 | 0 | 1 | 6 | 7 | 0 | 1 | 0 | 0 | 0 | 119 | 1 | 0 | 68 | 0 | 0 |
|  |  | 595 | 30 | 2 | 0 | 2 | 1 | 0 | 0 | 0 | 4 | 0 | 0 | 0 | 146 | 6 | 0 | 402 | 2 | 0 |
|  |  | 675 | 18 | 3 | 0 | 3 | 5 | 2 | 3 | 0 | 4 | 0 | 0 | 5 | 70 | 15 | 0 | 539 | 1 | 7 |
|  | **Total** | 3,736 | 257 | 23 | 2 | 16 | 14 | 91 | 31 | 25 | 20 | 10 | 0 | 6 | 1,044 | 32 | 2 | 2,141 | 13 | 9 |
|  | **Mean** | 311.3 | 21.4 | 1.9 | 0.2 | 1.3 | 1.2 | 7.6 | 2.6 | 2.1 | 1.7 | 0.8 | 0.0 | 0.5 | 87.0 | 2.7 | 0.2 | 178.4 | 1.1 | 0.8 |

**Table S4.**
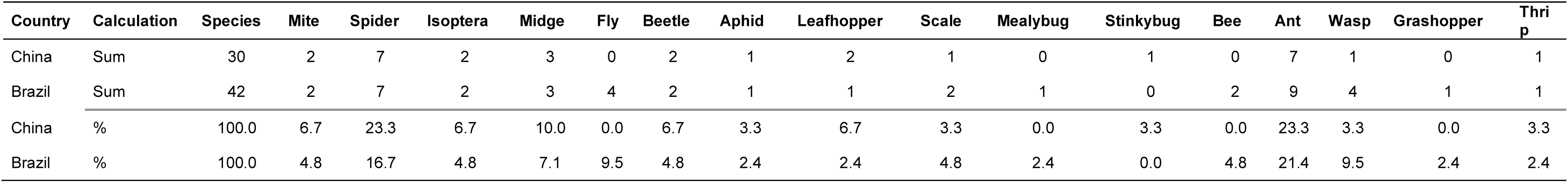
Summary of cocoa flower visitor morpho-species sorted by taxonomic groups recorded with ECV in our 21 research plots from Hainan, China and Bahia, Brazil.

| Country | Calculation | Species | Mite | Spider | Isoptera | Midge | Fly | Beetle | Aphid | Leafhopper | Scale | Mealybug | Stinkybug | Bee | Ant | Wasp | Grashopper | Thrip |
| --- | --- | --- | --- | --- | --- | --- | --- | --- | --- | --- | --- | --- | --- | --- | --- | --- | --- | --- |
| China | Sum | 30 | 2 | 7 | 2 | 3 | 0 | 2 | 1 | 2 | 1 | 0 | 1 | 0 | 7 | 1 | 0 | 1 |
| Brazil | Sum | 42 | 2 | 7 | 2 | 3 | 4 | 2 | 1 | 1 | 2 | 1 | 0 | 2 | 9 | 4 | 1 | 1 |
| China | % | 100.0 | 6.7 | 23.3 | 6.7 | 10.0 | 0.0 | 6.7 | 3.3 | 6.7 | 3.3 | 0.0 | 3.3 | 0.0 | 23.3 | 3.3 | 0.0 | 3.3 |
| Brazil | % | 100.0 | 4.8 | 16.7 | 4.8 | 7.1 | 9.5 | 4.8 | 2.4 | 2.4 | 4.8 | 2.4 | 0.0 | 4.8 | 21.4 | 9.5 | 2.4 | 2.4 |

**Table S5.** Classification criteria used to assign cocoa flower visitors to three basic ecological functions and their combinations. An example of classified images is shown in Supplementary Figure S3.

| Function | Description | Morphospecies ID |  |
| --- | --- | --- | --- |
|  |  | China | Brazil |
| Forager | The morphospecies visits reproductive structures (staminodes, style, and/or anthers), visibly feeding on nectar and/or pollen. Pollen is sometimes present on the body. | • cn_midge_01, 02<br>• cn_ant_02, 05 | • br_ant_03, 05, 07, 08<br>• br_bee_01, 02<br>• br_fly_01, 02, 03, 04<br>• br_midge_01, 02 |
| Herbivore | The morphospecies does not visit reproductive parts. Instead, it remains on petals with visible evidence of piercing, chewing, or cutting floral tissues. | • cn_beetle_01<br>• cn_leafhopper_01, 02<br>• cn_scale_01<br>• cn_ant_06 | • br_ant_01<br>• br_beetle_02<br>• br_grasshopper_01<br>• br_leafhopper_01<br>• br_mealybug_01<br>• br_scale_01, 02 |
| Predator | The morphospecies does not visit reproductive parts. It shows clear predatory behavior such as scouting for prey, chasing, or feeding on other arthropods. | • cn_spider_01, 02, 03, 04, 05, 06, 07 | • br_spider_01, 02, 03, 04, 05, 06, 07 |
| Forager/herbivore | The morphospecies shows evidence of both foraging and herbivory behaviors. | • cn_aphid_01<br>• cn_thrip_01 | • br_aphid_01<br>• br_beetle_01<br>• br_thrip_01 |
| Forager/predator | The morphospecies shows evidence of both foraging and predatory behaviors. | • cn_mite_02<br>• cn_wasp_01 | • br_mite_01, 02<br>• br_wasp_01 |
| Herbivore/predator | The morphospecies shows evidence of both herbivory and predatory behaviors. |  | • br_ant_04 |
| Not identified (NI) | The morphospecies remains stationary or exhibits non-specific movement on the flower, or short visit, without showing any visible foraging, herbivory, or predatory behavior. | • cn_beetle_02<br>• cn_isoptera_01, 02<br>• cn_stinkybug_01<br>• cn_ant_01, 02, 03, 04, 07<br>• cn_midge_01 | • br_ant_02, 06, 09<br>• br_isoptera_01, 02<br>• br_midge_03<br>• br_wasp_02, 03, 04 |

**Table S6.**
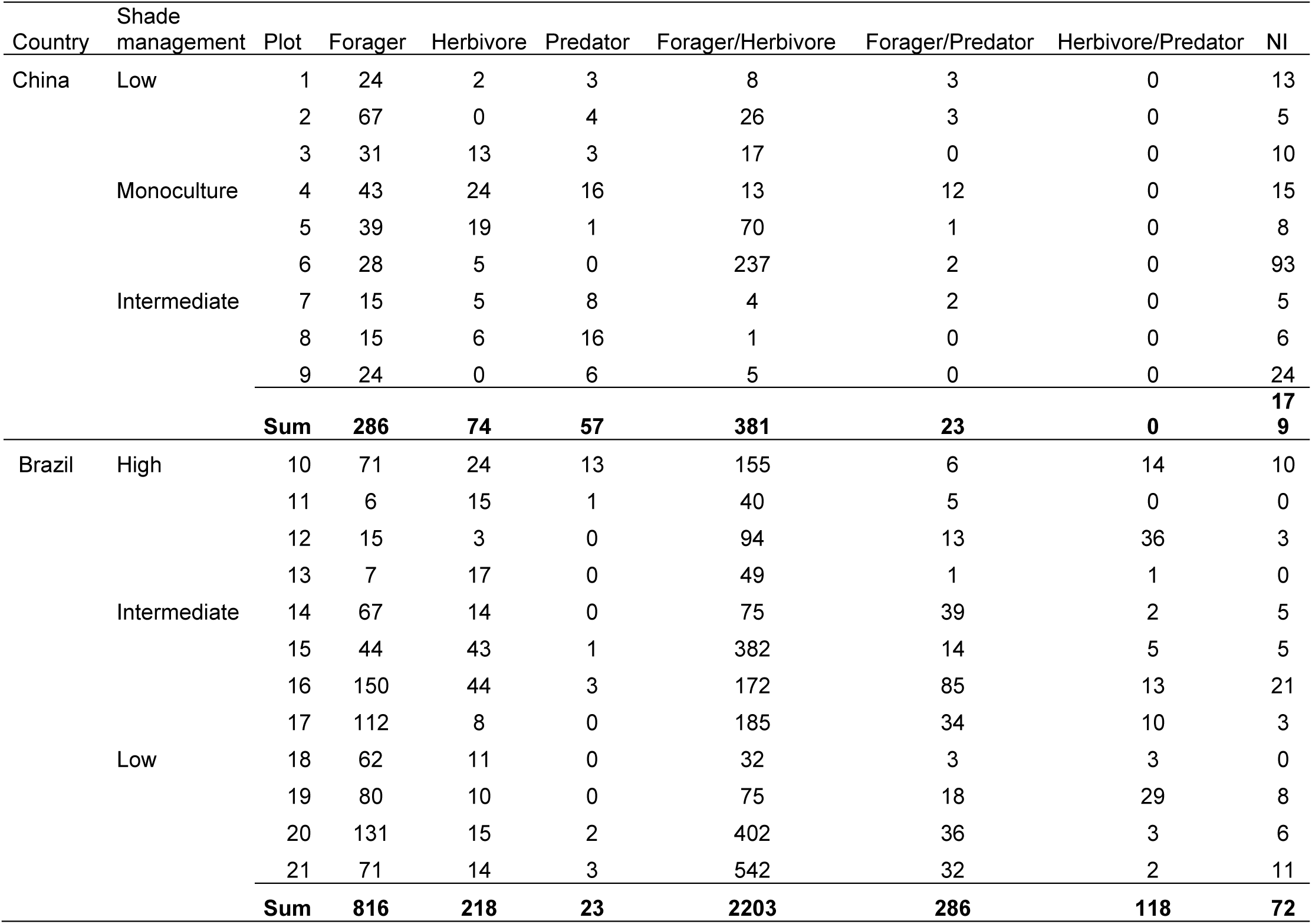
Summary of visits sorted by the three basic ecological functions and their combinations recorded with ECV in cocoa systems of Hainan, China and Bahia, Brazil.

**Table S7.**
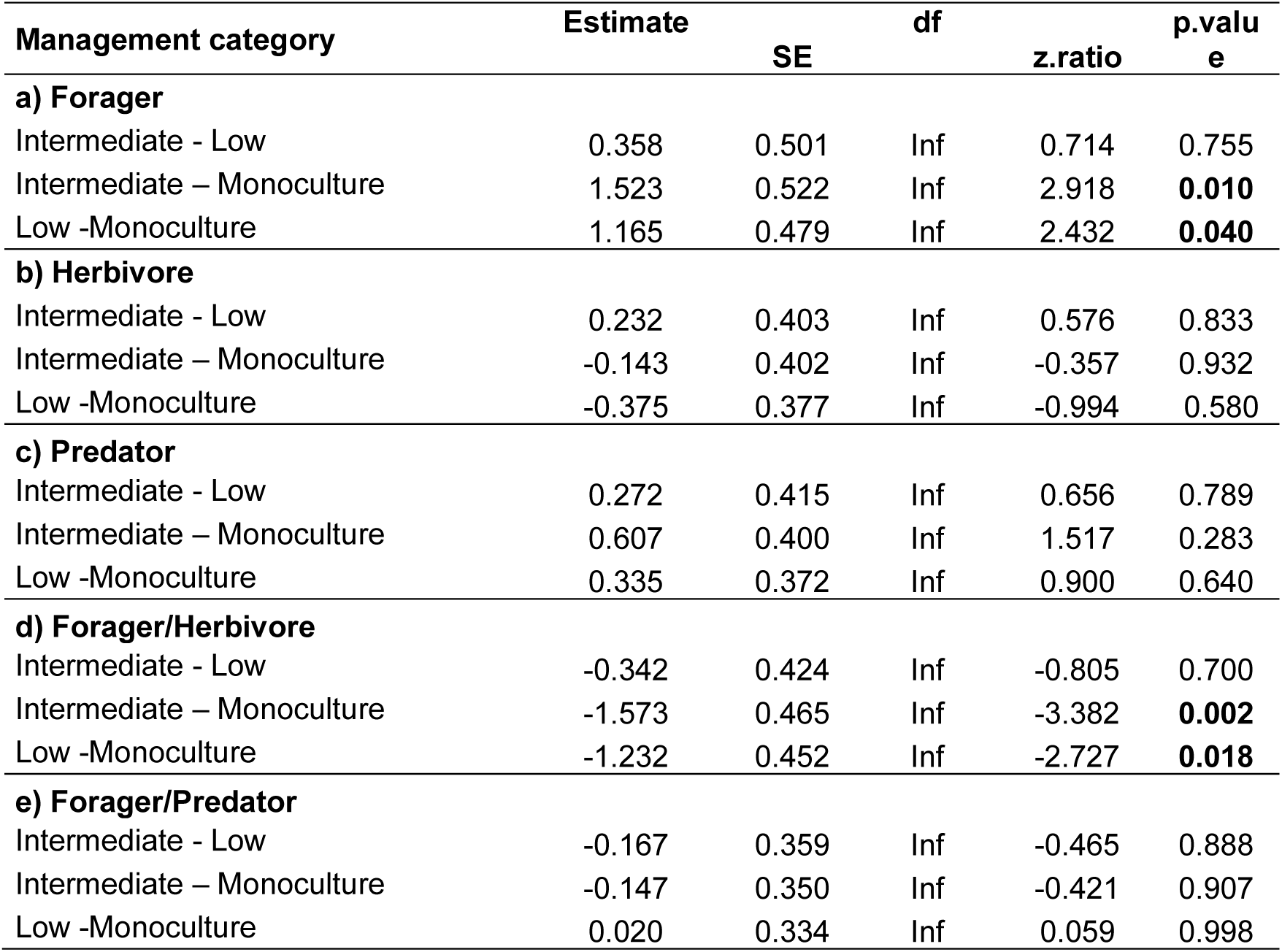
Summary results of the post-hoc multiple comparison of means analysis on shade-tree species richness and canopy cover effect on cocoa flower visitor ecological functions in Hainan, China. We split our nine plots in three system categories based on the number of shade-tree species and percentage of canopy cover. Categories are monocultures (i.e. no shade trees), low (i.e. one areca palm species and <30% canopy cover), and intermediate (i.e one coconut palm species and 40-50% canopy cover). Significant (p<0.05) effects are in bold.

**Table S8.**
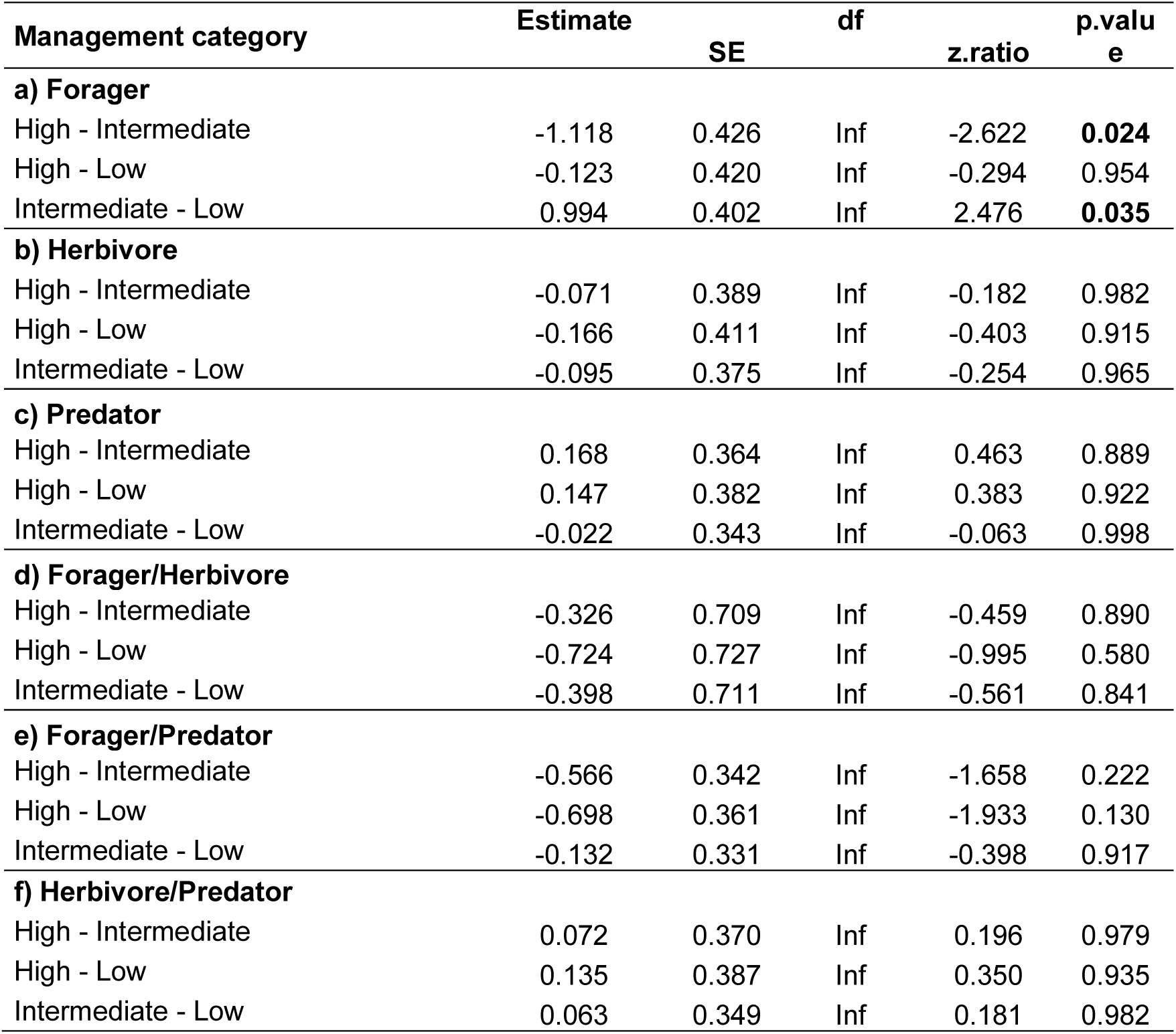
Summary results of the multiple comparison analysis on shade-tree species richness and canopy cover effect on cocoa flower visitor ecological functions in Brazil. We split our 12 plots in three system categories based on the number of shade-tree species and percentage of canopy cover. Categories are high (i.e. 7-8 shade-tree species and >50% canopy cover), intermediate (i.e. 4-5 shade-tree species and 40-50% canopy cover), and low (i.e. 1-3 shade-tree species and 30-40% canopy cover). Significant (p<0.05) effects are in bold.

**Table S9.** The coefficient estimates of our piecewise structural equation models (SEMs) where we investigated the interactions between temperature, canopy cover, flowering, fruit set, and main flower visitors (i.e. aphid, ant, midge) in cocoa systems of Hainan, China. Std.Error represents standard error, df is degree of freedom, Crit.Value is critical value, P.Value is p-value, and Std.Estimate is standard estimate. Significant p-values (<0.05) are in bold.

| Response | Predictor | Estimate | Std.Error | df | Crit.Value | P.Value | Std.Estimate |
| --- | --- | --- | --- | --- | --- | --- | --- |
| Flowering | Temperature | 0.069 | 0.040 | 207 | 1.746 | 0.082 | 0.127 |
| Flowering | Canopy cover | -0.002 | 0.006 | 207 | -0.312 | 0.756 | -0.024 |
| Flowering | Aphid | -0.108 | 0.102 | 207 | -1.056 | 0.292 | -0.081 |
| Flowering | Ant | 0.030 | 0.102 | 207 | 0.298 | 0.766 | 0.021 |
| Flowering | Midge | -0.113 | 0.092 | 207 | -1.230 | 0.220 | -0.088 |
| Fruit set | Temperature | 0.006 | 0.011 | 206 | 0.506 | 0.613 | 0.037 |
| Fruit set | Canopy cover | 0.001 | 0.002 | 206 | 0.666 | 0.506 | 0.050 |
| Fruit set | Flowering | 0.027 | 0.020 | 206 | 1.407 | 0.161 | 0.097 |
| Fruit set | Aphid | 0.041 | 0.029 | 206 | 1.418 | 0.158 | 0.108 |
| Fruit set | Ant | 0.021 | 0.029 | 206 | 0.725 | 0.469 | 0.051 |
| Fruit set | Midge | 0.070 | 0.026 | 206 | 2.722 | **0.007** | 0.195 |
| Aphid | Temperature | -0.051 | 0.027 | 208 | -1.920 | 0.056 | -0.126 |
| Aphid | Canopy cover | -0.018 | 0.004 | 208 | -4.416 | **<0.001** | -0.290 |
| Aphid | Ant | 0.117 | 0.069 | 208 | 1.710 | 0.089 | 0.109 |
| Aphid | Midge | -0.186 | 0.061 | 208 | -3.058 | **0.003** | -0.195 |
| Ant | Temperature | -0.001 | 0.027 | 209 | -0.038 | 0.970 | -0.003 |
| Ant | Canopy cover | -0.004 | 0.004 | 209 | -0.963 | 0.337 | -0.069 |
| Ant | Midge | -0.177 | 0.060 | 209 | -2.938 | **0.004** | -0.199 |
| Midge | Temperature | -0.017 | 0.031 | 210 | -0.563 | 0.574 | -0.041 |
| Midge | Canopy cover | 0.006 | 0.005 | 210 | 1.325 | 0.187 | 0.096 |

**Table S10.** The coefficient estimates of our piecewise structural equation models (SEMs) where we investigated the interactions between temperature, canopy cover, flowering, fruit set, and main flower visitors (i.e. thrip, ant, mite, midge) in cocoa systems of Bahia, Brazil. Std.Error represents standard error, df is degree of freedom, Crit.Value is critical value, P.Value is p-value, and Std.Estimate is standard estimate. Significant p-values (<0.05) are in bold.

| Response | Predictor | Estimate | Std.Error | df | Crit.Value | P.Value | Std.Estimate |
| --- | --- | --- | --- | --- | --- | --- | --- |
| Flowering | Temperature | -0.109 | 0.038 | 255 | -2.868 | **0.005** | -0.174 |
| Flowering | Canopy cover | -0.003 | 0.004 | 255 | -0.924 | 0.356 | -0.056 |
| Flowering | Thrip | 0.121 | 0.028 | 255 | 4.355 | **<0.001** | 0.266 |
| Flowering | Ant | 0.016 | 0.037 | 255 | 0.420 | 0.675 | 0.026 |
| Flowering | Midge | 0.502 | 0.784 | 255 | 0.640 | 0.523 | 0.038 |
| Flowering | Mite | 0.029 | 0.061 | 255 | 0.474 | 0.636 | 0.028 |
| Fruit set | Temperature | -0.003 | 0.013 | 254 | -0.262 | 0.794 | -0.016 |
| Fruit set | Canopy cover | -0.003 | 0.001 | 254 | -2.810 | **0.005** | -0.169 |
| Fruit set | Flowering | -0.044 | 0.020 | 254 | -2.154 | **0.032** | -0.134 |
| Fruit set | Thrip | -0.005 | 0.009 | 254 | -0.510 | 0.611 | -0.032 |
| Fruit set | Ant | 0.048 | 0.012 | 254 | 4.045 | **<0.001** | 0.245 |
| Fruit set | Midge | -0.034 | 0.254 | 254 | -0.135 | 0.893 | -0.008 |
| Fruit set | Mite | 0.007 | 0.020 | 254 | 0.356 | 0.722 | 0.021 |
| Thrip | Temperature | -0.096 | 0.085 | 256 | -1.128 | 0.261 | -0.070 |
| Thrip | Canopy cover | 0.011 | 0.008 | 256 | 1.344 | 0.180 | 0.083 |
| Thrip | Ant | -0.272 | 0.081 | 256 | -3.359 | **0.001** | -0.204 |
| Thrip | Midge | -0.502 | 1.761 | 256 | -0.285 | 0.776 | -0.017 |
| Thrip | Mite | -0.207 | 0.137 | 256 | -1.509 | 0.133 | -0.092 |
| Ant | Temperature | 0.003 | 0.066 | 257 | 0.039 | 0.969 | 0.003 |
| Ant | Canopy cover | -0.009 | 0.006 | 257 | -1.416 | 0.158 | -0.089 |
| Ant | Midge | -0.821 | 1.356 | 257 | -0.605 | 0.546 | -0.038 |
| Ant | Mite | 0.079 | 0.105 | 257 | 0.750 | 0.454 | 0.047 |
| Midge | Temperature | 0.004 | 0.003 | 258 | 1.407 | 0.161 | 0.089 |
| Midge | Canopy cover | 0.000 | 0.000 | 258 | -0.331 | 0.741 | -0.021 |
| Midge | Mite | -0.001 | 0.005 | 258 | -0.202 | 0.840 | -0.013 |
| Mite | Temperature | -0.054 | 0.038 | 259 | -1.392 | 0.165 | -0.087 |
| Mite | Canopy cover | -0.001 | 0.004 | 259 | -0.259 | 0.796 | -0.016 |

**Table S11.**
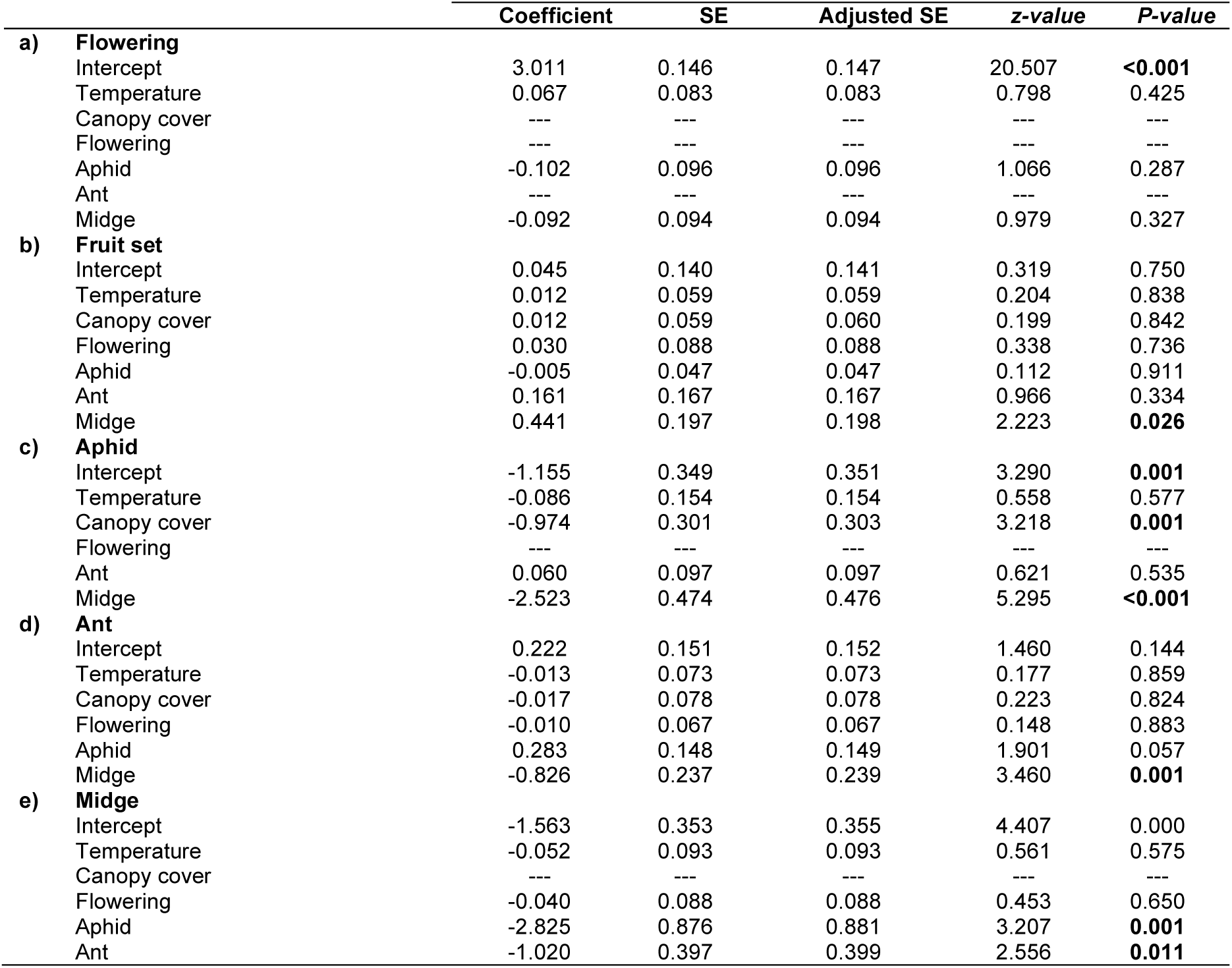
Generalized linear mixed effect (GLMMs) model selection outputs of flowering (a), fruit set (b), and main flower visitors (c-e) drivers in cocoa systems of Hainan, China. We used glmmTMB package (Brooks et al., 2023) to test the effect of i) temperature, ii) canopy cover, and iii) main flower visitors (i.e. aphid, ant, midge) on i) flowering, ii) fruit set, and iii) main flower visitors. We fitted our models with a negative binomial distribution, except for fruit set fitted with a binomial distribution to account for overdispersion in the binary data. We included plot as a random effect. We then followed a model selection approach to identify the most parsimonious models (ΔAICc≤ 2) and averaged the models coefficients using the “model.avg” function from the “MuMIn” package (Barton, 2009). Significant p-values (>0.05) are in bold. We then contrasted these results with our SEM models (Tabel S9) to identify consistency in our results.

**Table S12.** Generalized linear mixed effect (GLMMs) model selection outputs of flowering (a), fruit set (b), and main flower visitors (c-f) drivers in cocoa systems of Bahia, Brazil. We used glmmTMB package (Brooks et al., 2023) to test the effect of i) temperature, ii) canopy cover, and iii) main flower visitors (i.e. thrip, ant, mite, midge) on i) flowering, ii) fruit set, and iii) main flower visitors. We fitted our models with a negative binomial distribution, except for fruit set fitted with a binomial distribution to account for overdispersion in the binary data. We included plot as a random effect. We then followed a model selection approach to identify the most parsimonious models (ΔAICc≤ 2) and averaged the models coefficients using the “model.avg” function from the “MuMIn” package (Barton, 2009). Significant p-values (>0.05) in bold. We then contrasted these results with our SEM models (Table S10) to identify consistency in our results.

|  | Coefficient | SE | Adjusted SE | z-value | P-value |
| --- | --- | --- | --- | --- | --- |
| **a) Flowering** |  |  |  |  |  |
| Intercept | 2.931 | 0.186 | 0.187 | 15.686 | **<0.001** |
| Temperature | -0.133 | 0.058 | 0.058 | 2.306 | **0.021** |
| Canopy cover | --- | --- | --- | --- | --- |
| Thrip | --- | --- | --- | --- | --- |
| Ant | 0.206 | 0.056 | 0.056 | 3.664 | **<0.001** |
| Mite | 0.013 | 0.034 | 0.034 | 0.397 | 0.691 |
| Midge | 0.009 | 0.027 | 0.027 | 0.320 | 0.749 |
| **b) Fruit set** |  |  |  |  |  |
| Intercept | -2.048 | 0.248 | 0.249 | 8.213 | **<0.001** |
| Temperature | -0.010 | 0.078 | 0.078 | 0.129 | 0.897 |
| Canopy cover | -0.373 | 0.229 | 0.230 | 1.625 | 0.104 |
| Flowering | -0.736 | 0.349 | 0.351 | 2.096 | **0.036** |
| Thrip | -0.042 | 0.145 | 0.146 | 0.289 | 0.772 |
| Ant | 0.367 | 0.153 | 0.154 | 2.383 | **0.017** |
| Mite | 0.012 | 0.080 | 0.081 | 0.151 | 0.880 |
| Midge | --- | --- | --- | --- | --- |
| **c) Thrip** |  |  |  |  |  |
| Intercept | 1.938 | 0.161 | 0.162 | 11.949 | **<0.001** |
| Temperature | --- | --- | --- | --- | --- |
| Canopy cover | 0.048 | 0.112 | 0.113 | 0.429 | 0.668 |
| Flowering | 0.312 | 0.132 | 0.133 | 2.346 | **0.019** |
| Ant | -0.032 | 0.072 | 0.072 | 0.442 | 0.659 |
| Mite | -0.079 | 0.138 | 0.138 | 0.573 | 0.567 |
| Midge | --- | --- | --- | --- | --- |
| **d) Ant** |  |  |  |  |  |
| Intercept | 1.351 | 0.125 | 0.126 | 10.758 | **<0.001** |
| Temperature | -0.031 | 0.088 | 0.088 | 0.354 | 0.723 |
| Canopy cover | -0.048 | 0.096 | 0.096 | 0.495 | 0.621 |
| Flowering | --- | --- | --- | --- | --- |
| Thrip | -0.221 | 0.138 | 0.138 | 1.603 | 0.109 |
| Mite | 0.003 | 0.035 | 0.036 | 0.079 | 0.937 |
| Midge | -0.024 | 0.080 | 0.080 | 0.295 | 0.768 |
| **e) Mite** |  |  |  |  |  |
| Intercept | -0.067 | 0.252 | 0.253 | 0.266 | 0.790 |
| Temperature | -0.035 | 0.139 | 0.139 | 0.250 | 0.803 |
| Canopy cover | -0.026 | 0.128 | 0.129 | 0.202 | 0.840 |
| Flowering | 0.132 | 0.236 | 0.237 | 0.556 | 0.578 |
| Thrip | -0.111 | 0.222 | 0.222 | 0.500 | 0.617 |
| Ant | 0.015 | 0.107 | 0.107 | 0.138 | 0.890 |
| Midge | -0.007 | 0.076 | 0.077 | 0.096 | 0.924 |
| **e) Midge** |  |  |  |  |  |
| Intercept | -2.808 | 0.282 | 0.283 | 9.920 | **<0.001** |
| Temperature | 0.042 | 0.148 | 0.148 | 0.284 | 0.776 |
| Canopy cover | -0.014 | 0.109 | 0.110 | 0.123 | 0.902 |
| Flowering | 0.020 | 0.104 | 0.104 | 0.188 | 0.851 |
| Thrip | --- | --- | --- | --- | --- |
| Ant | -0.038 | 0.189 | 0.190 | 0.199 | 0.843 |
| Mite | -0.018 | 0.166 | 0.167 | 0.106 | 0.915 |

